# A major locus controls a biologically active pheromone component in *Heliconius melpomene*

**DOI:** 10.1101/739037

**Authors:** Kelsey J.R.P. Byers, Kathy Darragh, Jamie Musgrove, Diana Abondano Almeida, Sylvia Fernanda Garza, Ian A. Warren, Pasi M. Rastas, Marek Kučka, Yingguang Frank Chan, Richard M. Merrill, Stefan Schulz, W. Owen McMillan, Chris D. Jiggins

## Abstract

Understanding the production, response, and genetics of signals used in mate choice can inform our understanding of the evolution of both intraspecific mate choice and reproductive isolation. Sex pheromones are important for courtship and mate choice in many insects, but we know relatively little of their role in butterflies. The butterfly *Heliconius melpomene* uses a complex blend of wing androconial compounds during courtship. Electroantennography in *H. melpomene* and its close relative *H. cydno* showed that responses to androconial extracts were not species-specific. Females of both species responded equally strongly to extracts of both species, suggesting conservation of peripheral nervous system elements across the two species. Individual blend components provoked little to no response, with the exception of octadecanal, a major component of the *H. melpomene* blend. Supplementing octadecanal on the wings of octadecanal-rich *H. melpomene* males led to an increase in the time until mating, demonstrating the bioactivity of octadecanal in *Heliconius.* Using quantitative trait locus (QTL) mapping, we identified a single locus on chromosome 20 responsible for 41% of the parental species’ difference in octadecanal production. This QTL does not overlap with any of the major wing color or mate choice loci, nor does it overlap with known regions of elevated or reduced *F*_ST_. A set of 16 candidate fatty acid biosynthesis genes lies underneath the QTL. Pheromones in *Heliconius* carry information relevant for mate choice and are under simple genetic control, suggesting they could be important during speciation.

## Introduction

Chemical communication is the oldest form of sensory communication, and plays a fundamental role in the ecology of organisms across the tree of life. In terms of reproductive behavior, chemical communication is involved in premating isolation in a variety of systems from orchids (Peakall et al. 2010) to *Drosophila* (Shahandeh et al. 2017) to cichlids (Plenderleith et al. 2005), highlighting its importance as a mediator of speciation and diversification (Smadja & Butlin 2009). Of particular interest is the evolution of pheromones, chemical compounds that mediate intraspecies interactions. In particular, signaling (production and emission of pheromone compounds) and receiving (reception and interpretation of chemical signals) components are predicted to evolve in concert to maintain reproductive isolation between closely related species in sympatry (Smadja & Butlin 2009). Despite its ubiquity, chemical communication has been less well studied than *e.g.* visual communication. The result is that outside of a limited set of (relatively well-studied) examples we know relatively little about the role that chemical signals play in reproductive isolation and evolution. However, recent technical advances (e.g. later-generation gas chromatography-coupled mass spectrometry and gas chromatography-coupled electroantennographic detection) mean that chemical communication is increasingly accessible for evolutionary studies.

Studies of pheromones have been most widespread in insects, in particular in the order Lepidoptera and in *Drosophila* (Smadja & Butlin 2009). In Lepidoptera, work on pheromones has largely focused on long-range female pheromones of nocturnal moths (often due to their economic importance), where pheromone divergence commonly contributes to speciation and relatively simple structural variations in pheromones are known to produce drastic differences in mate attraction. For example, different populations of the corn borer moth *Ostrinia nubialis* exhibit a simple *cis* to *trans* switch in the pheromone 11-tetradecenyl acetate (Kochansky et al. 1975, Lassance et al. 2010) that leads to partial reproductive isolation (Dopman et al. 2010). Sex pheromones have been less well studied in day-flying butterflies, where visual signaling is often assumed to play a more dominant role in mate choice (Vane-Wright and Boppré 1993, Löfstedt et al. 2016). However, male butterflies also emit close-range pheromone bouquets which may act in concert with other wing pattern and behavioral cues (Mérot et al. 2015), and are important in mate choice (Darragh et al. 2017) as well as decreasing heterospecific mating (Mérot et al. 2015).

Despite the potential importance of pheromones in butterflies, aphrodisiac pheromones have so far been identified in only eight butterfly species (Meinwald et al. 1969; Pliske and Eisner 1969; Grula et al. 1980; Nishida et al. 1996; Schulz and Nishida 1996; Andersson et al. 2007; Nieberding et al. 2008; Yildizhan et al. 2009). This is in stark contrast to the approximately two thousand species of moths where female pheromones or attractants are known (Löfstedt et al. 2016). There is a similar absence of knowledge about the genetic basis of variation in pheromone production (but see Liénard et al. 2014).

As in other diurnal butterflies, the chemical bouquets of the genus *Heliconius* (Darragh et al. 2017; Mann et al. 2017), are complex, both in identity and quantity of compounds, though just a few individual compounds may be biologically active pheromones. Although studying variation in pheromones within and across species can point to potential candidates (e.g. Darragh et al. 2019b), identifying which components of these complex chemical bouquets are responsible for pheromonal communication is a considerable challenge. This is particularly true as even minor compounds can have major effects (McCormick et al. 2014; Chen et al. 2018). Determining pheromone bioactivity requires screening compounds via physiological activity followed by behavioral verification, and in many cases pheromone bouquet composition is known but the bioactive components remain unidentified. In addition, behavioral outcomes may differ despite similar responses in the peripheral nervous system (Chen & Fadamiro 2007; Seeholzer et al. 2018).

Here we take advantage of two closely related *Heliconius* butterfly species, *H. melpomene* and *H. cydno*, to further our knowledge of the ecology, evolution and genetics of male lepidopteran pheromones. The two species diverged about 2.1 million years ago (Arias et al 2014; Kozak et al. 2015), and are strongly reproductively isolated (Jiggins 2017). Over the past decade there has been considerable research into the genomic architecture of differences in wing pattern and male mate preference (Jiggins 2017; Merrill et al. 2019). Surprisingly, both wing color and male mating preferences between these species have a relatively simple genetic basis with a large proportion of the difference between parental forms being controlled by a small handful of loci of large effect (Naisbit 2003; Jiggins 2017; Merrill et al. 2019). This has important implications for speciation, as theory predicts that large effect loci contribute to speciation in the face of gene flow (Via 2012). Similarly, tight physical linkage between loci that contribute to isolating barriers will facilitate speciation (Felsenstein 1981; Merrill et al. 2010; Smadja and Butlin 2011), and there is evidence for tight linkage of a gene controlling wing pattern (*optix*) and a major effect QTL underlying divergent male preference behaviors (Merrill et al. 2019). Pheromonal differences between the species might also be expected to be under control by major effect loci, as has been seen in a variety of moth species (Groot et al. 2016, Haynes et al. 2016). The extent to which loci underlying pheromone production overlap with wing pattern and mate choice loci is unclear, but given the existing linkage between wing pattern and male mate choice loci, we might predict additional linkage of pheromone production with wing pattern loci.

Recent research has demonstrated the importance of male pheromones in mating success (Darragh et al. 2017). To better characterize male butterfly pheromone production and the role it plays in mating and species recognition, we comprehensively analyzed pheromone bouquets of *Heliconius melpomene* and *H. cydno* butterflies to determine the most important bioactive compounds. We carried out electrophysiology and behavioral experiments and identified octadecanal as a biologically active pheromone component. To better understand the genetic basis of octadecanal production and determine the location of loci responsible for the production of octadecanal relative to loci involved in wing color pattern and male mating preference, we mapped loci responsible for differences in the level of octadecanal synthesis between the two species. We found that a single locus on chromosome 20 explains 41% of the parental species differences in this pheromone, with no linkage between this locus and known color pattern and male mate choice loci.

## Materials and Methods

Data and analysis code are deposited in Dryad under accession XXX. Sequencing reads leading to linkage map construction are deposited in the European Nucleotide Archive (ENA) under project PRJEB34160.

### Butterflies

Stocks from central Panama of *Heliconius melpomene rosina* and *H. cydno chioneus* (hereafter *H. melpomene* and *H. cydno)* were used for all experiments (Darragh et al. 2017). Butterflies were reared in insectaries at the Smithsonian Tropical Research Institute, Gamboa, Panama under ambient temperature and light conditions. Eggs were collected from these breeding stocks and the resulting larvae fed on *Passiflora platyloba var. williamsi, P. biflora, P. menispermifolia*, and *P. vitifolia* until pupation. Data from (Darragh et al. 2019a) indicate that larval diet does not affect the major compounds found in *H. melpomene*, suggesting that this dietary variation is unlikely to affect results. Newly-eclosed adult butterflies were separated by sex to ensure virgin status and supplied with flowers from *Psiguria warscewiczii, Psiguria triphylla, Gurania eriantha, Psychotria poeppigiana, Stachytarpheta mutabilis*, and *Lantana* sp. (most likely *L. camara)* as pollen sources, as well as a ~20% sucrose solution. All experiments used virgin butterflies. For assessment of *H. cydno* wing bouquets, male butterflies were between 10-12 days post eclosion and had fed exclusively on *Passiflora platyloba var. williamsi.* For electrophysiology, female butterflies were between 1 and 20 days post eclosion, and males between 10-20 days post eclosion to ensure male sexual maturity (Darragh et al. 2017). For behavior, female butterflies were used the day after eclosion and males were between 10 and 59 days post eclosion. Natural wing extracts of both species were extracted from males 10-12 days post eclosion as described in (Darragh et al. 2017) using dichloromethane plus 1 ng/μL 2-tetradecyl acetate (hereafter “DCM+IS”) and concentrated approximately 10x prior to use under still room air. All samples were stored at −20°C before use.

### Identification and quantification of androconial compounds

To identify species-specific compounds among our two species, the chemical composition of the *H. cydno* androconial bouquet was investigated in samples from 26 adult male *H. cydno* and compared with 31 adult male *H. melpomene*, the latter including samples previously analyzed in (Darragh et al. 2017; Darragh et al. 2019a), all collected as above. Samples were assessed using gas chromatography-mass spectrometry (GC-MS) with an Agilent 7890B gas chromatograph coupled with an Agilent 5977 mass spectrometer with electron ionization (Agilent Technologies, California, USA). The GC utilized an Agilent HP-5MS capillary column (30m length, 0.25mm inner diameter), helium carrier gas at 1.2 mL/minute, and an Agilent ALS 7693 autosampler. Injection was splitless with an inlet temperature of 250°C. The temperature ramp was isothermal at 50°C for 5 minutes, then increased at 5°C/minute to 320°C and was then isothermal for 5 minutes. Samples were identified using a custom MS library and quantified by comparison with the internal standard.

In line with (Darragh et al. 2017), wings from eight male *H. cydno* were dissected into four regions: hindwing androconia, forewing overlap region, hindwing rest-of-wing, and forewing rest-of-wing, and all extracted identically after dissection. Wing region area was quantified by photographing the wings before dissection and measuring the total pixel area of each wing region in the GNU Image Manipulation Program v.2.8.20 (GIMP Development Team), with the pixel-mm conversion via measurement of a ruler in the photograph. Quantified compounds in each wing region for each individual were scaled by the area of the relevant wing region in that individual.

### Chemicals

Syringaldehyde (3,5-dimethoxy-4-hydroxybenzaldehyde), 1-octadecanol, and henicosane were obtained commercially (Sigma-Aldrich). The aldehydes octadecanal, (Z)-11-icosenal, and (Z)-13-docosenal were obtained from the respective alcohols 1-octadecanol, (Z)-11-icosen-1-ol, and (Z)-13-docosen-1-ol by oxidation with iodoxybenzoic acid (IBX) in ethyl acetate according to (More and Finney 2002). The required alcohols (Z)-11-icosen-1-ol and (Z)-13-docosen-1-ol were in turn obtained by lithium aluminum hydride reduction from commercially available (Z)-11-icosenoic acid and methyl (Z)-13-docosenoate (Larodan) according to (Cha and Brown 1993). The seven target compounds (see Figures S4 and S5 for structures and reaction scheme) were chosen due to their quantitative dominance in the chemical profiles of *H. melpomene* and *H. cydno.* The solvent for all synthesized compounds was hexane, with the exception of the polar syringaldehyde, which was diluted in a 1:10 mixture of dichloromethane and hexane.

Synthetic blends of *H. melpomene* and *H. cydno* male wing bouquets were prepared from these synthesized compounds. Synthetic *H. melpomene* contained 23.2 ng/μL syringaldehyde, 23.3 ng/μL octadecanal, 6.9 ng/μL 1-octadecanol, 4.7 ng/μL (Z)-11-icosenal, 20.3 ng/μL (Z)-11-icosenol, and 4.8 ng/μL (Z)-13-docosenal. Synthetic *H. cydno* contained 47.0 ng/μL syringaldehyde and 93.3 ng/μL henicosane. Floral direct extractions from *Lantana* sp. (most likely *L. camara)* growing wild in Gamboa, Panama were used as a positive control. Single umbels were removed from plants at dawn and placed in a scintillation vial to which 400μL of DCM+IS was added. After 1 hour, the DCM+IS was removed to a glass vial and kept at −20°C before use.

### Electroantennography

Electrophysiological preparations were assembled as follows: antennae were cut from the head of a virgin butterfly using fine scissors and the final 6.5 segments (approximately 1.5mm, avoiding cutting at the segment boundary) cut off with a scalpel. Both antennae were then placed in parallel across an antenna fork (Syntech, Buchenbach, Germany) and held in place using electrode gel (Spectra 360 Electrode Gel, Parker Laboratories Inc., Fairfield, NJ, USA). The antenna fork was mounted on a Syntech EAG CombiProbe with 10x internal gain, and signal from this was routed through a Syntech IDAC4. EAG waveforms recorded using Syntech GcEad/2014 software. Stimulus pulses were delivered using a Syntech CS-55 Stimulus Controller with foot pedal trigger. Both continuous unhumified room air and stimulus pulses were delivered to the preparation at 1.5 liters/min through a tube of approximately 8mm inner diameter. The stimulus pulses were delivered in triplets of 0.5 seconds each, separated by 5 seconds, with triplets initiated every 30 seconds. Stimulus delivery used odor cartridges assembled from Pasteur pipettes with a strip of filter paper plugged with cotton when not in use; each stimulus cartridge was ‘charged’ with 10uL of stimulus solution for each experiment. Each antennal preparation was used only once.

Two sets of stimuli were delivered: a species comparison set and a synthetic compound set. Both sets used air (nothing added to filter paper), hexane, and DCM+IS as negative controls and *Lontono* extract as a positive control. The species comparison set included male wing extracts from *H. melpomene* and *H. cydno* (“Mnat” and “Cnat” respectively) and synthetic blends representing the two species (“Msyn” and Csyn”). The synthetic compound set included air, hexane, DCM+IS, the conspecific male wing extract, the conspecific synthetic blend, and the seven synthetic compounds. Presentation order was randomized before each experiment. Species comparison experiments consisted of sixteen pulses each of the seven stimuli, interspersed with five pulses of *Lantana* extract at the start, between every three stimuli, and at the end. Synthetic compounds were similar, with eleven pulses of each of the twelve stimuli, interspersed with four pulses of *Lontono* at the start, between every four stimuli, and at the end. For analysis, the first triplet of each stimulus set was removed, leaving fifteen and ten pulses respectively. At least ten female and five male butterflies of each species were used with each experiment.

Onset time and amplitude of EAG responses (i.e. the magnitude of the decrease from the baseline signal, barred lines in Figure S6) were marked using GcEad/2014. To control for antennal signal degradation over time, a time-dependent correction factor was calculated using linear interpolation between each *Lontono* set and this applied to the absolute value of the EAG response amplitude. These corrected amplitudes were then scaled to the amplitude of the initial *Lontono* set to partially control for differences between preparations.

### Analysis of antennal adaptation

Short-term adaptation (STA), as defined in (Zufall and Leinders-Zufall 2000), is adaptation occurring only over a very brief window from the initial stimulus that then resolves quickly with no further stimulation (e.g. the 10 second interval given for salamanders in the reference). By contrast, long-term or long-lasting adaptation (LTA) is defined in the same source as persisting over an extended period of time, up to several minutes in vertebrates and insects (Stengl 2010), with recovery of response upon presentation of a different stimulus. Within our electrophysiological data set, there is the potential to measure both types of adaptation; since we corrected for preparation degradation over time, we should be able to measure true LTA and STA. STA, if present, should be evident within an individual triplet, as the stimuli within a triplet are separated by 5 seconds (and thus STA is likely to persist within a triplet), whereas LTA should be evident across an individual stimulus set, as these lasted 5.5-8 minutes with maximum intervals of 30 seconds between stimuli, insufficient for LTA to be abolished if we assume similar mechanisms as vertebrates. We assessed antennal responses for LTA by pooling all triplets within a stimulus-species-sex combination. We pooled these data within stimulus set types (species comparison and synthetic compound), treating butterfly preparation identity as a random effect. For each stimulus, the change in antennal response was assessed over the time since initial presentation of the stimulus. STA was assessed by looking for significant changes in the residuals from this analysis between members of a triplet.

### Behavior

To test the potential role of octadecanal in *H. melpomene* female mate choice, behavioral experiments were conducted in insectaries at STRI, Gamboa, Panama, between April and July 2018. One day old virgin females were presented with both an octadecanal treated and a control *H. melpomene* male for two hours. Males were at least ten days old and were selected based on similarity of size, wing-wear and age, with treatment being allocated randomly by coin flip. Either 25 μl octadecanal solution (140 ng/μl octadecanal in hexane, thus adding 3500ng to the existing average 773.4ng for approximately a 5.5x dose) or 25 μl pure hexane (both evaporated down to a smaller volume of approximately 10 μl under room air) was applied to the hindwing androconial region of each male, and males were then allowed 30 minutes to settle before beginning the two hour experiment period. Experiments began at or close to 9am, with observations being made every 15 minutes or until mating occurred. *Heliconius melpomene* was chosen for these experiments as an adequate number of individuals were available, although the less easily reared (and thus less available) *H. cydno* might have provided a clearer picture of the role of octadecanal in premating reproductive isolation.

To test the persistence of the octadecanal treatment on the wings of live butterflies, a separate set of *H. melpomene* males was treated as above with either hexane or octadecanal. Separate males were sampled at 30 minutes post treatment and two hours post treatment by extraction of the forewing overlap region (Darragh et al. 2017) and the hindwing androconia in DCM+IS as above, with two males per treatment-time combination. Octadecanal was then measured using GC-MS as above and quantified by peak area comparison with the 2-tetradecyl acetate internal standard.

### Quantitative trait locus mapping for octadecanal production

To map the genetic basis for octadecanal production in *H. melpomene*, we took advantage of the fact that *H. cydno* produces little to no octadecanal. Bidirectional F_1_ crosses between the two species revealed that the *H. cydno* phenotype (low to no octadecanal) is dominant over the high octadecanal production found in *H. melpomene*, so we constructed backcross families by crossing F_1_ males to female *H. melpomene* from our existing stocks. A total of ten families (nine with a female *H. melpomene* grandparent and one with a female *H. cydno* grandparent) were constructed, with each offspring representing a single recombination event from the F_1_ father. We constructed backcrosses to *H. cydno* (127 individuals from 15 families) in addition, as some segregation was seen in this backcross direction as well. Butterflies were reared and wing extracts collected and analyzed from male offspring as described above, except that all larvae were reared on *Passiflora platyloba var. williamsi.* Bodies of male offspring were collected into dimethyl sulfoxide (DMSO) for later library preparation. The Castle-Wright estimators for octadecanal and octadecanol production were calculated using the phenotypic variance of the backcross individuals as the estimated segregation variance (Jones 2001).

Qiagen DNeasy kits (Qiagen) were used for DNA extraction. Individuals were genotyped either by RAD-sequencing as previously described (Davey et al. 2017; Merrill et al. 2019), or low-coverage whole genome sequencing using nextera-based libraries. For the nextera-based libraries a secondary purification was performed using magnetic SpeedBeads™ (Sigma) dissolved in 0.44mM PEG8000, 2.5M NaCl, 1mM Tris-Cl pH=8, and 0.1mM EDTA pH=8.0. High-throughput sequencing libraries were generated using a method based on the Nextera DNA Library Prep (Illumina, Inc.) with purified Tn5 transposase (Picelli et al. 2014). Sample barcoding was performed using PCR extension with an i7-index primer (N701-N783) and the N501 i5-index primer. Libraries were purified and size selected using the same beads as above. Pooled libraries were sequenced by HiSeq 3000 (Illumina) by BGI (China).

Linkage mapping was conducted using standard Lep-MAP3(LM3) pipeline (Rastas 2017). First, individual fastq files were mapped to the melpomene reference genome using BWA MEM (Li 2011) and then sorted bams were created using SAMtools (Li and Durbin 2011). The input genotype likelihoods were constructed by SAMtools mpileup and pileupParser2+pileup2posterior from LM3. The pedigree of individuals was validated and corrected using IBD (identity-by-descent) module and the sex of individuals was validated and corrected according to the coverage on the Z chromosome and autosomes using SAMtools depth. Then, ParentCall2 (parameter ZLimit=2) and Filtering2 (dataTolerance=0.001) modules were called on the input data and a random subset of 25% of markers (to speed up analysis) was used for the subsequent steps.

Initial linkage groups (chrX.map) and marker orders (orderX.txt) were constructed based on the *H. melpomene* genome for each of 21 chromosomes. SeparateChromosomes2 was run on each of these groups with the default parameters (lodLimit=10) except for map=chrX.map (for X=1..21). Finally, OrderMarkers2 was run on each chromosome in the constructed order, with parameter scale=0.05, recombination2=0, evaluateOrder=orderX.txt and map=result_from_SeparateChromsomes2.chrX.txt. Another evaluation was done with data in the grandparental phase (additional parameter grandparentPhase=1). The phased data of these orders were matched using phasematch script (LM3) and obtaining all markers from the first evaluation in the grandparental phase. This obtained result was used as the final map.

Map construction resulted in the retention of 447,818 SNP markers across 89 and 127 individuals with phenotype data in backcrosses to *H. melpomene* and *H. cydno* respectively. To facilitate computation, markers were thinned evenly by a factor of ten, resulting in 44,782 markers with no missing data. Octadecanal production was log-transformed to obtain normality, then regressed against marker position using the R/qtl2 R library (Broman et al. 2018). Significance thresholds were obtained by permutation testing in R/qtl2 with 1000 permutations, and QTL confidence intervals obtained using the bayesjnt command. To account for the family structure present in our QTL mapping populations, we additionally included a kinship matrix calculated by R/qtl2 using the LOCO (leave one chromosome out) method in the marker regression and recalculated significance thresholds and confidence intervals with the kinship term included. Percent variance explained was calculated as the difference in phenotype means of individuals of each genotype divided by the difference in the parental phenotype. Since the genetic linkage map was based on whole genome data, we were able to obtain physical positions of QTL confidence interval endpoints. The physical positions of the kinship-included confidence interval were used to query Lepbase (Challis et al. 2016) for potential candidate genes from the *H. melpomene* genome. To identify putative functions for each potential candidate, protein sequences from Lepbase were searched against the nr (non-redundant) protein database using BLASTp (Altschul et al. 1990). For each candidate with a promising functional annotation, exons were pulled out of the *H. cydno* genome (Pessoa Pinharanda 2017) and aligned to the *H. melpomene* genes using BLASTn with each *H. melpomene* exon to search for SNPs between the two species. Selection was tested at the sequence level using codeml (Yang 2007) in pairwise mode with the F3X4 codon frequency model.

### Statistical analysis

Differences in individual compounds between *H. melpomene* and *H. cydno* were assessed using a Welch’s *t*-test. Wing region differences in *H. cydno* were assessed for each individual compound found in at least four of eight samples of at least one wing region using a linear mixed model with wing region as a fixed effect and butterfly identity as a random effect using the package nlme v.3.1.137 (Pinheiro et al. 2018). Statistical tests for these models were assessed with the Anova function in the package car v.3.0.0 (Fox and Weisberg 2011), and comparisons between wing regions were performed using the emmeans package v.1.3.1 (Lenth 2018).

For electroantennography, species comparison sets and synthetic compound sets were analyzed separately. Corrected and scaled EAG responses (for each experiment within sexes and species) were compared between stimuli with a linear mixed model with stimulus as a fixed effect, butterfly preparation identity as a random effect, and the interaction of stimulus and preparation as a random effect using nlme as above. Statistical tests for these models were assessed with the Anova and emmeans (version 1.1.3) functions as above.

Long-term adaptation was assessed using robust linear mixed models with the package robustlmm v.2.2.1 (Koller 2016), with corrected but unscaled amplitude as the response variable, time since initial presentation of the stimulus as a fixed variable, and butterfly preparation as a random variable. Responses were pooled across samples within a species-sex combination and considered for each stimulus separately. As robustlmm does not provide p-values, confidence intervals were used to assess significance and difference between sample-species combinations. Short-term adaptation was assessed using the residuals from the same regression. Differences in the residuals between triplets within a triplet set were tested using a one-sample t-test with a hypothesized mean value of zero (i.e. no difference between residuals), performed on the subtractive difference between the residuals of the third (last) and first triplets.

Female mate choice was assessed using a binomial test, and treatment differences in time until mating were assessed with a two-sided t-test assuming unequal variances. Octadecanal persistence was not assessed statistically due to the small sample size. All statistical tests were performed in R v.3.5.0, v.3.5.1, or v.3.5.2 (R Core Team 2013).

## Results

### Androconial chemical bouquets of *Heliconius melpomene* and *H. cydno*

In order to identify candidate pheromone compounds, we first investigated the distribution of chemical compounds on the wings of *H. cydno* for comparison with published data on *H. melpomene* (Figure 1). The chemical profile of the wings of the two species were quite different, with few shared major compounds. We found that the bouquet of *H. cydno* was simpler than that of *H. melpomene*, with 7 main compounds versus 21 in *H. melpomene. Heliconius cydno* also had a less abundant overall bouquet, with an individual total of 1787 ± 776 ng vs. 3174 ± 1040 ng in *H. melpomene* (Table S1). Most of the main compounds (defined as those occurring in at least 90% of individuals) in *H. cydno* were linear alkanes (4 of 7 compounds), while *H. melpomene* has a more diverse array of different compound classes. Of the five major compounds (> 100ng per individual) in *H. melpomene*, only syringaldehyde was found in similar amounts in *H. cydno*;, the major compounds octadecanal, 1-octadecanol, (Z)-11-icosenal, and (Z)-11-icosenol were absent or found in very low amounts in *H. cydno.* Comparison with previously published data for other *Heliconius* species, all of which lack octadecanal in the large amounts seen in *H. melpomene*, demonstrates that this high level of octadecanal is an evolutionarily derived state in *H. melpomene* (Mann et al. 2017; Darragh et al. 2019b). When focusing on the hindwing androconia of *H. cydno*, only two compounds [syringaldehyde (24.7% of the hindwing androconial bouquet) and (Z)-11-icosenol (1.7%)] were specific to this region. For details, please see SI text, Figures S1-S3, and Tables S1-S2.

**Figure 1:**
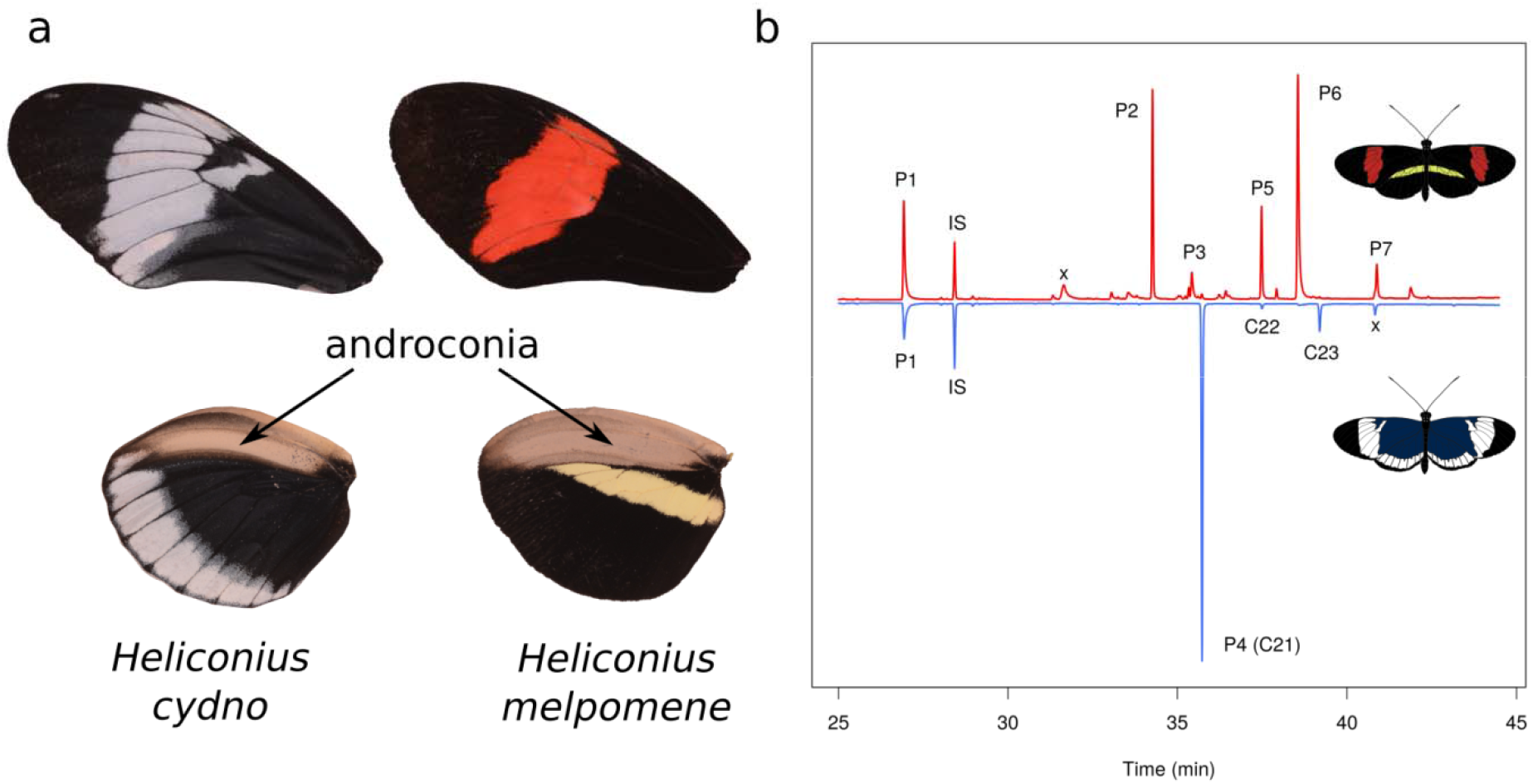
Androconial chemistry of *Heliconius melpomene* and *H. cydno.* a: dorsal forewing and hindwing of each species showing the silvery androconial region of the hindwing used during male courtship, b: Total ion chromatogram of *H. melpomene* (top) and *H. cydno* (bottom) wing androconia. P1: syringaldehyde; P2: octadecanal; P3: 1-octadecanol; P4: henicosane; P5: (Z)-11-icosenal; P6: (Z)-11-icosenol; P7: (Z)-13-docosenal; IS: internal standard (2-tetradecyl acetate); x: contaminant; C21: henicosane; C22: docosane; C23: tricosane.

### Electroantennograŋhic responses to con- and heterospecific pheromone stimulus sets

We next investigated the electroantennographic (EAG) response of both species to natural con- and heterospecific pheromone bouquets extracted from adult male butterflies. In general, EAG responses were more pronounced in females than in males, and we did not see a pattern of increased response to conspecific pheromone bouquets over those of heterospecifics. Females of both *H. melpomene* and *H. cydno* responded more strongly (i.e. showed a larger voltage displacement from the antennal baseline) to both natural wing extracts (Mnat and Cnat, respectively) as compared with the control solvent (DCM+IS) (Figure 2, Figure S6; see Table S3 for statistical details). Males of *H. melpomene* also responded to both wing extracts, while no response was seen in male *H. cydno*, likely due to large inter-individual variation in response in this species-sex combination. Females and males of both species showed equivalent responses to *H. melpomene* and *H. cydno* wing extracts.

**Figure 2:**
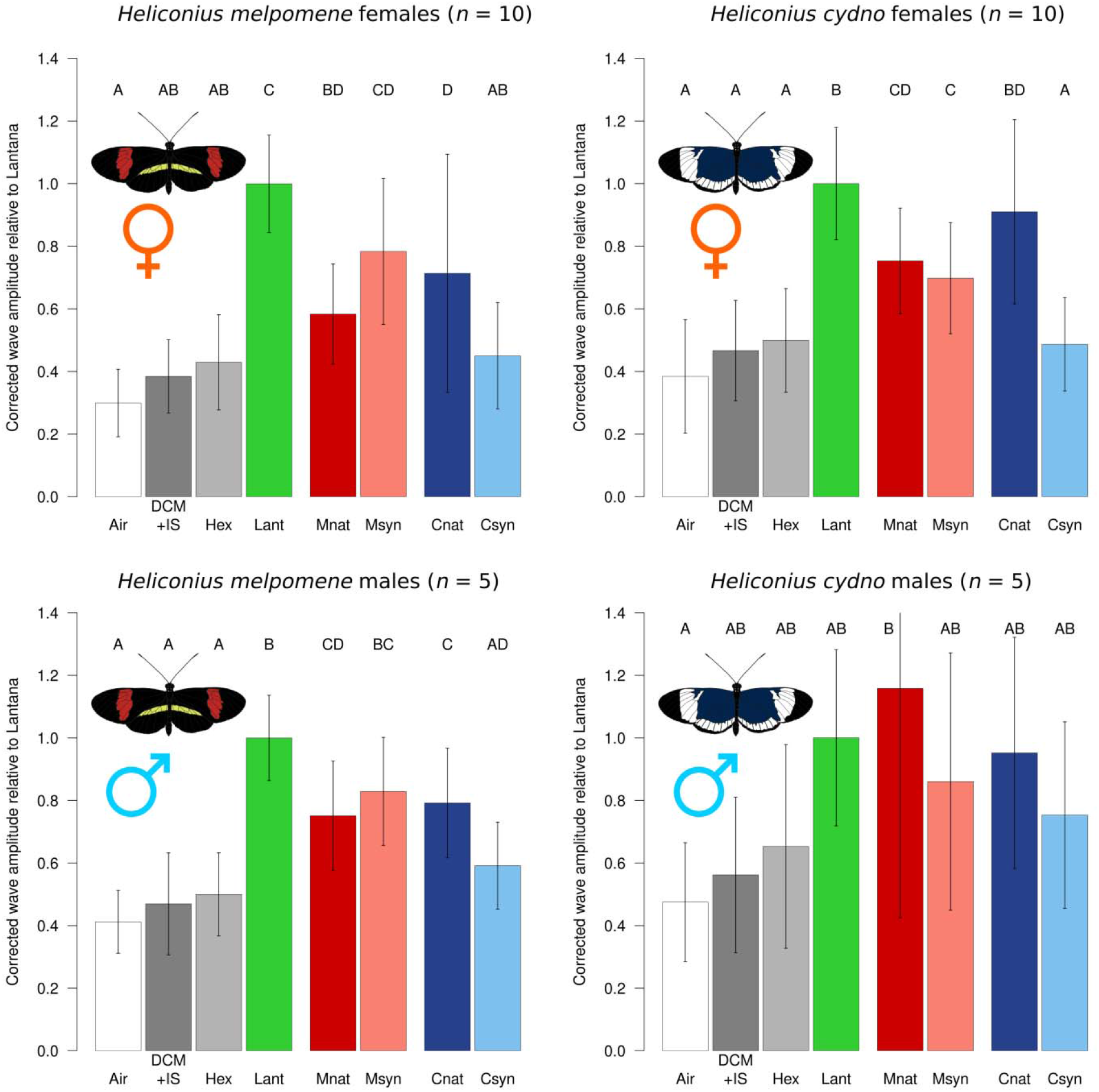
Electroantennographic responses of *Heliconius* butterflies to conspecific and heterospecific wing extracts. Stimuli: air (white), dichloromethane plus 2-tetradecyl acetate (internal standard) (dark gray), hexane (light gray), *Lantana* extract (green), natural male *H. melpomene* wing extract (red), synthetic *H. melpomene* blend (pink), natural male *H. cydno* wing extract (dark blue), synthetic *H. cydno* blend (blue). Bars: average of normalized corrected amplitude ± standard deviation.

We then explored antennal responses to synthetic compound blends. These were based on the most abundant compounds from each species (see Methods, Figure S4). We were able to successfully recapitulate the pheromone of *H. melpomene*, but not that of *H. cydno.* Male and female *H. melpomene* responded equally to the natural *H. melpomene* wing extract and its synthetic wing blend (Msyn) in both stimulus sets (Figures 2 and 3). By contrast, both sexes of *H. cydno* evidenced no increased response to the synthetic *H. cydno* wing blend (Csyn) when compared with the hexane solvent. In all cases this response was lower than to natural *H. cydno* wing extract, indicating that we have not successfully identified its active component(s).

**Figure 3:**
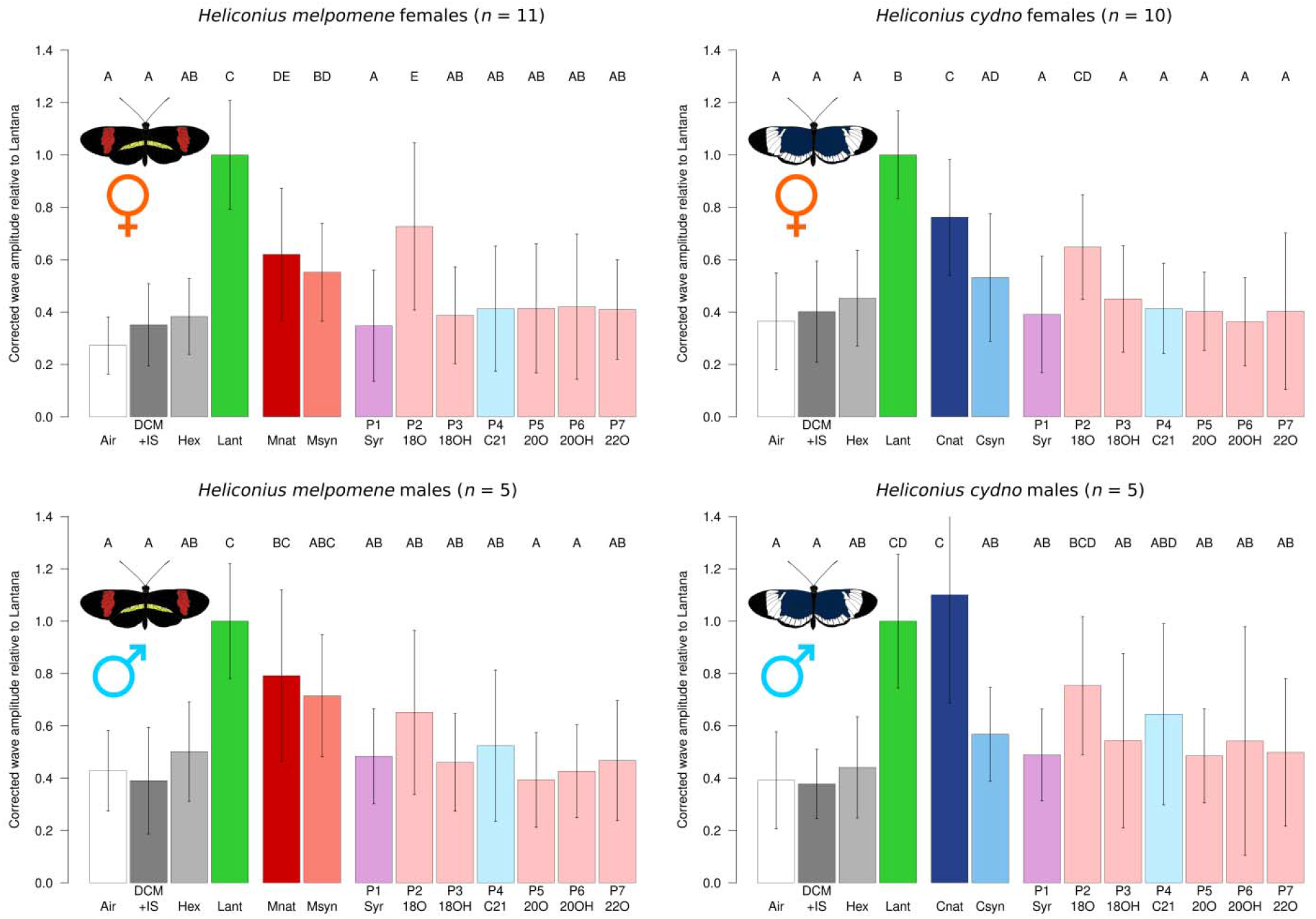
Electroantennographic responses of *Heliconius* butterflies to synthetic wing compounds. Stimuli: air (white), dichloromethane plus 2-tetradecyl acetate (internal standard) (dark gray), hexane (light gray), *Lantana* extract (green), natural male *H. melpomene* or *H. cydno* wing extract (red or dark blue respectively), synthetic *H. melpomene* or *H. cydno* blend (pink or blue respectively). PI (Syr): syringaldehyde; P2 (180): octadecanal; P3 (180H): 1-octadecanol; P4 (C21): henicosane; P5 (200): (Z)-11-icosenal; P6 (200H): (Z)-11-icosenol; P7 (220): (Z)-13-docosenal. Bars: average of normalized corrected amplitude ± standard deviation. Light pink: compound part of synthetic *H. melpomene* blend; light blue: compound part of synthetic *H. cydno* blend; purple: compound in both *H. melpomene* and *H. cydno* synthetic blends.

### Electroantennographic responses to individual pheromone components from both species

Finally, we explored the responses to individual compounds to identify specific biologically active pheromone components. Only octadecanal differed significantly from the controls in any species-sex combination (Figure 3, Table S3), and this difference was seen only in females. As our experiments used concentrations approximately 10x those present in nature (approximately equivalent to the concentrated natural extracts, except (Z)-13-docosenal which was at 30x based on prior chemical analysis), this is unlikely to be due to differences in compound abundance in our experiments. No other compound was significantly different from hexane. In female *H. melpomene*, response to octadecanal was stronger than response to the *H. melpomene* synthetic mixture, suggesting a slight inhibitory response due to the presence of other synthetic compounds in the mixture, though no single compound produced this inhibition in isolation. By contrast, male *H. melpomene* responded equally to the conspecific synthetic mixture and octadecanal (as well as the solvent hexane), and both female and male *H. cydno* responded equally to their conspecific synthetic mixture and both its components (syringaldehyde and henicosane), with no evidence for a synergistic mixture effect. Antennal responses to a given stimulus can change over time, and this may reflect biological processes of neuronal adaptation. Female *H. melpomene* adapted equally quickly to octadecanal and the natural and synthetic *H. melpomene* pheromones, while adaptation to other stimuli was equal to the control, further supporting octadecana?s salience as the main pheromone in *H. melpomene* (see SI text, Figures S7-S8, and Table S4).

### Behavioral response to octadecanal supplementation in *H. melpomene*

We next confirmed a behavioral response to the most physiologically active substance, octadecanal, one of the dominant compounds in *H. melpomene.* A total of 29 behavioral trials were conducted in *H. melpomene*, with mating observed in 18 (62%); one trial was excluded due to wing damage, leaving 17 successful matings. With our small sample size, we found no evidence that females showed a preference for either treatment, mating with the hexane male 11 times (65%) and with the octadecanal male the remaining 6 times (35%) (p = 0.332). However, mating latency (time from experiment onset to mating) was significantly longer for the octadecanal matings than for the hexane matings (average 88.5 versus 43.7 minutes; t = 2.7848, df = 8.2491, p = 0.023). There was no evidence that this mating latency was due to evaporation of the octadecanal treatment, as there was no detectable drop in octadecanal quantity in the hindwing androconia over the duration of the experiment [comparison of 30 minute and 2 hour treatments (Figure S9)], although some octadecanal was lost initially before the experiment began. Furthermore, little octadecanal rubbed off onto the forewing overlap region. Interestingly, although about 5.5x as much octadecanal as normal should have been present on the wings of treated males, only about 2-2.5x was seen after 30 minutes (the time at which the female would be introduced to the two males), suggesting some of the added octadecanal was lost before the start of the behavioral experiments, perhaps due to oxidization or pheromone hydrolysis, as has been shown in some moths (Ferkovich et al. 1982).

### Genetic basis of octadecanal production in *H. melpomene*

Analysis of octadecanal production by Fı males showed that the *H. cydno* octadecanal phenotype (little to no octadecanal) is dominant over the octadecanal-rich *H. melpomene* phenotype, and octadecanal production segregates in backcrosses to *H. melpomene* (Figure S10). Using the variance within the *H. melpomene* backcross individuals, we calculated a Castle-Wright estimator of 0.81 loci, suggesting a potentially monogenic basis for octadecanal production in *H. melpomene.* Quantitative trait locus (QTL) mapping with 89 individuals from 10 families revealed a single significant peak on chromosome 20 (Figure 4a, Figure S11). The chromosome 20 peak remained significant when kinship was taken into account and explained 41.31% of the difference between the two parent species. Bayesian confidence intervals for the peak on chromosome 20 were identical with and without kinship, spanning a range of 46.9-56.37cM with the peak at 47.66cM (with or without kinship), corresponding to a physical range of 3.4Mb along chromosome 20. To ensure that our findings were replicable across individual families, we also constructed effect plots at the kinship peak for each family separately, and all showed the same directionality (Figure 4b). The peak on chromosome 20 does not overlap with any of the major wing color loci (Jiggins 2017; Van Belleghem et al. 2017), nor does it overlap with mate choice QTL (Merrill et al. 2019). The confidence interval region contains 160 genes, all of which represent potential candidates for octadecanal production. Although octadecanal production appeared recessive in F_1_ individuals, there was also some segregation seen in backcrosses to *H. cydno*, so we performed QTL mapping on these additional individuals (127 individuals, 15 families). This analysis recapitulated the peak on chromosome 20 with a similar confidence interval (42.35-54.85cM with a peak at 45.76cM), providing independent support for this QTL peak from an entirely different set of hybrid individuals.

**Figure 4:**
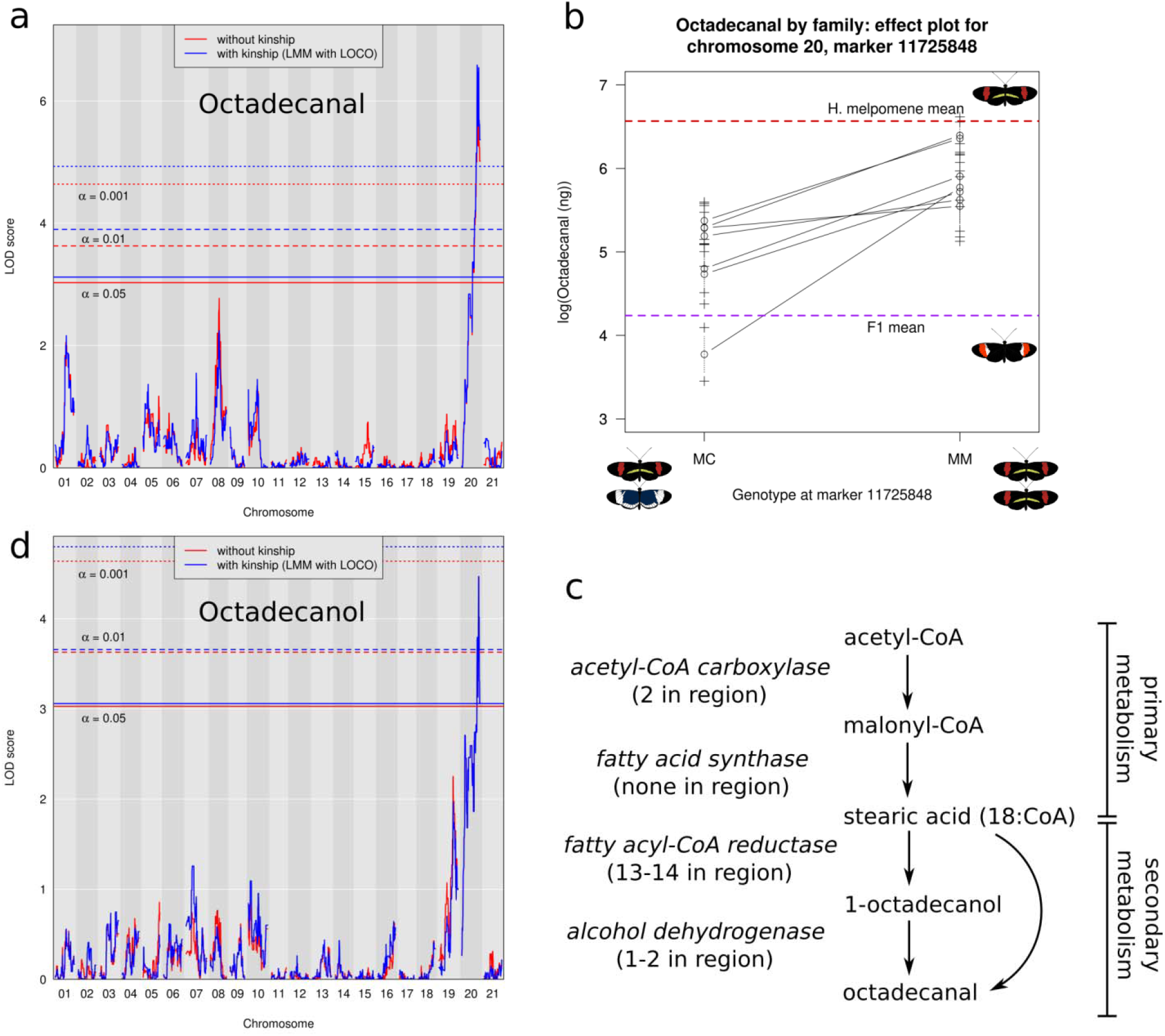
QTL mapping of octadecanal and octadecanol production in *Heliconius melpomene.* a: QTL map for production of octadecanal. b: Effect plots at the peak of the locus on chromosome 20 for the seven individual backcross mapping families with at least 5 individuals, c: Potential biosynthetic pathway for octadecanal production, d: QTL map for production of octadecanol, a potential precursor of octadecanal.

We next evaluated the evidence in support of the 160 genes in this interval to identify top candidates for octadecanal production. In total, 14 were putative fatty-acyl CoA reductases (FARs), which catalyze the conversion of fatty acids to alcohols via a bound aldehyde intermediate (Table S5). Octadecanal is most likely produced via this pathway (Figure 4c), either as a direct product of a FAR-catalyzed conversion of 18-carbon stearic acid (by releasing the bound intermediate directly) or as a product of a further dehydrogenation of the alcohol intermediate (octadecanol) to the aldehyde product. Two candidate alcohol dehydrogenases, which might catalyze this reaction, were also contained within the region, yielding a total of 16 candidates. To ascertain whether octadecanol might serve as the precursor to octadecanal in *H. melpomene*, we also searched for QTL underlying octadecanol production (Castle-Wright estimator of 0.71 loci), and found a very similar pattern to octadecanal, with a single QTL peak on chromosome 20 (Figure 4d) explaining 25.36% of the difference between the two parent species.

This peak broadly overlapped the octadecanal peak, with a much broader confidence interval from 10.91-56.37cm (12.9Mb) and a peak at 51.82cM regardless of whether kinship was taken into account (Figure S11). The 14 FARs in the region are highly clustered, with a set of eight found within a 133kb region. Sequence comparison between the *H. melpomene* and *H. cydno* alleles showed that nearly all of these genes harbor nonsynonymous SNPs between the two species (Table S5, Supplementary Data 1). One gene showed no coding SNPs between the two species; nine had between three and 24 nonsynonymous SNPs; and four had more substantial changes, including missing exons and frameshifts. The final two genes (one alcohol dehydrogenase and one FAR) could not be found in the *H. cydno* genome, and may instead represent annotation or assembly errors in *H. melpomene* or, alternately, deletions in *H. cydno*. Nearly all of the intact genes displayed purifying selection (ω between 0.001 and 0.2459), with only the remaining alcohol dehydrogenase (ω = 1.2478) under positive selection (Table S5). Taken together, these results suggest that either of the alcohol dehydrogenase candidates may underlie the production of bioactive octadecanal from octadecanol in *H. melpomene*, although functional experiments are required to confirm this hypothesis. Alternately, as QTL for octadecanal and its likely precursor octadecanol overlap, a single FAR may be responsible for producing both volatiles.

## Discussion

Previous work has shown that male *Heliconius* butterflies use aphrodisiac pheromones during courtship, and the presence of these pheromones is necessary for successful mating to take place (Darragh et al. 2017). However, the identity of the bioactive pheromone components and the genetic basis underlying their production was unknown. Here we demonstrate that two closely related species with strong reproductive isolation, *H. melpomene* and *H. cydno*, show major differences in chemical bouquets (see also Mann et al. 2017; Darragh et al. 2019b). Strong divergence between closely related species is unusual in Lepidoptera. Instead, pheromone types are typically shared between closely related species, with only subtle differences in similar compounds or differences in ratios of the same compound (Löfstedt et al 2016). Somewhat surprisingly, despite these major differences in putative pheromone signals, we detected no difference in the strength of antennal response to wing chemical bouquets. Nonetheless, we have identified a single compound, octadecanal, which elicited a significant response in females of both species. Somewhat surprisingly, octadecanal was also physiologically active in *H. cydno* females, while it is largely absent from the male *H. cydno* wing bouquet.

These data on identical antennal responses may suggest that the peripheral nervous system of *H. melpomene* and *H. cydno* have not diverged in concert with their male wing chemistry. This is in contrast to similar published studies of other insect species. For example, the moth *Ostrinia nubialis*, whose *E*-and Z-strains diverged approximately 75,000-150,000 years ago (Malausa et al. 2007), strains have opposite topologies in the antennal lobe and antennal sensillae (Kárpáti et al. 2007; Koutroumpa et al. 2014). Similar divergence in peripheral nervous system architecture has been seen in *Rhagoletis pomonella* (Frey and Bush 1990; Tait et al. 2016) and *Drosophila mojavensis* (Date et al. 2013; Crowley-Gall et al. 2016) despite much shorter divergence times than between *H. melpomene* and *H. cydno* (approximately 2.1 million years ago, Arias et al 2014; Kozak et al. 2015).

The sensory periphery is only the first of many mechanisms that may influence mate choice, and it is increasingly clear that differences within the brain can be important in mate choice, even when detection mechanisms of the sensory periphery are conserved (Hoke et al. 2008; Hoke et al. 2010; Seeholzer et al. 2018). Our results in *Heliconius* are similar to those observed in *Colias* butterflies. The sister species *C. eurytheme* and *C. philodice* show very similar female electrophysiological responses to the con- and heterospecific pheromone compounds, despite a behavioral effect of treating males with heterospecific pheromones (Grula et al. 1980). *Colios* butterflies, like *Heliconius*, use multiple signals when choosing between mates (Papke et al. 2007), so rapid divergence in peripheral nervous system elements may not play a role in the evolution of reproductive isolation. In *Heliconius*, where EAG responses are very similar between species, differences in the antennal lobe and higher brain regions (e.g. the mushroom body or lateral protocerebrum, see e.g. Montgomery & Merrill 2017) may account for interspecies differences in mate choice behavior. Female *Heliconius* butterflies likely integrate multiple signals (including pheromones, male courtship flights, and visual cues) in these higher brain regions when making the decision to mate.

We have identified octadecanal as the major pheromone component in *H. melpomene* and showed that responses to it are conserved across both species despite its general absence in *H. cydno.* As a fully saturated unbranched compound, octadecanal is unusual in being unrelated to known female pheromone types in moths, which largely use unsaturated or methylated hydrocarbons with or without terminal functional groups (Löfstedt et al. 2016). The activity of octadecanal as a pheromone, however, has been tested behaviorally or electrophysiologically in eight species of Lepidoptera across a variety of families (Tatsuki et al. 1983; Cork et al. 1988; Tumlinson et al. 1989; Ho et al. 1996; McElfresh et al. 2000; Yildizhan et al. 2009; El-Sayed et al. 2011; Pires et al. 2015; Chen et al. 2018). Only in *Cerconota anonella* (Pires et al. 2015), and now in *Heliconius melpomene*, has some electrophysiological and behavioral activity been seen. The closely related hexadecanal is also a major pheromone component in the butterfly *Bicyclus anynana* (Nieberding et al. 2008), and differs from octadecanal only in its origin from palmitic rather than stearic acid and carbon number, so the role of octadecanal is not entirely unexpected.

Male pheromones categorized in other butterflies represent a wide range of chemical classes, including terpenoids (Meinwald et al. 1969; Pliske & Eisner 1969; Andersson et al. 2007), pyrrolizidine alkaloid derivatives (Meinwald et al. 1969; Pliske & Eisner 1969; Nishida et al. 1996; Schulz & Nishida 1996), macrolides (Yildizhan et al. 2009), aromatics (Andersson et al. 2003), fatty acid esters (Grula et al. 1980), and (in *Bicyclus anynana)* unsaturated fatty acid derived compounds more typical of moths (Nieberding et al. 2008). Male moth pheromones follow this wide distribution of chemical classes, with only a few species using fatty acid derived compounds, most notably *Heliothis virescens* which uses octadecanol, hexadecanol, and related compounds but not the respective aldehydes such as octadecanal (Conner & Iyengar 2016). In contrast to *H. melpomene*, we have failed to discover any physiologically active pheromones in *H. cydno*, perhaps because a minor component or components not tested here is biologically active (McCormick et al. 2014; Chen et al. 2018). Attempts to identify the *H. cydno* pheromone using GC-coupled electroantennographic detection were unfortunately unsuccessful due to technical issues with the setup, and thus the *H. cydno* pheromone remains undescribed.

Intriguingly, despite the strong EAG response, there was a marked negative behavioral response to increased octadecanal in *H. melpomene.* A plausible explanation for the negative behavioral response to octadecanal supplementation is that disruption of the normal mixture ratios of *H. melpomene* may inhibit the female response, as seen in the butterfly *Pieris napi*, where synergistic processing of two volatile components in the male bouquet is necessary for acceptance behavior (Larsdotter-Mellstrom et al. 2016). Octadecanal may also experience a dose-response curve with an aversive response to higher concentrations and an attractive response at lower ones. Potential mixture or dosage effects suggest that female *H. melpomene* may use octadecanal quantity or relative abundance to assess male quality or choose between courting males. The increased mating latency with octadecanal-treated males may reflect females undergoing a period of adjustment, either in the peripheral or central nervous system, to the higher dose of octadecanal; this would be consistent with our results showing long-term adaptation to octadecanal. We remain uncertain of what effect, if any, octadecanal would have on the behavior of *H. cydno*, where it is largely absent from the male pheromone bouquet, as we were unable to rear an adequate number of *H. cydno* for behavioral trials. It may be used to avoid courtship with *H. melpomene*, supporting other divergent signals such as color pattern in maintaining reproductive isolation between the two species (Jiggins et al. 2001).

Given the strong physiological and behavioral response to octadecanal, and its possible role in reproductive isolation between *H. melpomene* and *H. cydno*, we studied the genetic basis of differences between the two species. Fatty-acid derived compounds comprise the largest category of Lepidoptera sex pheromones (Ando and Yamakawa 2011), and are produced from fatty acyl-CoA precursors via the action of several enzymes. Since these pheromones are secondary metabolites derived from primary metabolic pathways, their production is likely to be relatively labile in evolutionary terms, allowing simple genetic changes to drive the wide diversity of lepidopteran sex pheromones. Even though we have a broad knowledge of pheromone diversity in Lepidoptera, our understanding of the genetics of pheromone biosynthesis is relatively weak. Pheromone gland-specific fatty acyl-CoA reductases have been identified in a number of moth species, although most are identified solely on transcriptomic analysis of the gland without functional characterization (Groot et al 2016, Löfstedt et al 2016). In the moth *Ostrinia* and butterfly *Bicyclus*, FARs involved in male pheromone biosynthesis have been identified and shown to use the same biosynthetic pathway as female pheromones (Lassance & Löfstedt 2009, Lienard et al 2014). Pheromone-producing alcohol oxidases, which potentially catalyze the conversion of antennally inactive octadecanol to the active component octadecanal, have not yet been described in any insect to our knowledge.

Using a QTL mapping approach, we have shown that the production of octadecanal has a relatively simple genetic basis, with a region on chromosome 20 corresponding to production of both octadecanal and its likely precursor octadecanol in *Heliconius.* This locus therefore likely represents a region under divergent selection between *H. melpomene* and *H. cydno* that is unlinked to previously identified species differences in color and mate choice (Jiggins 2017; Merrill et al. 2019). Patterns of F_ST_ between the species are highly heterogeneous and were not especially informative in further delimiting the locus (data from Martin et al. 2013). Due to our small mapping population, the confidence intervals for these QTL therefore remain large: the octadecanal QTL spans 3.4Mb and contains 160 genes. Of these, we identified 16 likely candidate genes based on known biosynthetic pathways in moths and the butterfly *Bicyclus anynana* (Liénard et al. 2014): 14 fatty acyl-CoA reductases and two alcohol dehydrogenases. Fatty acyl-CoA reductases have previously been identified in *H. melpomene* by (Liénard et al. 2014), who noted lineage-specific duplications within *H. melpomene* on two scaffolds corresponding to *H. melpomene* chromosomes 19 and 20. All but one of the candidate FARs found on chromosome 20 were identified by Liénard et al., but all fall outside their clade of pheromone gland FARs. The identified *Bicyclus* FAR that produces hexadecanol does not also produce the major pheromone hexadecanal, implying the presence of an additional as yet undescribed alcohol dehydrogenase in *Bicyclus.* The biochemical similarity between hexadecanal and octadecanal suggests *Heliconius*, like *Bicyclus*, may also use an alcohol dehydrogenase to produce octadecanal. By contrast, the overlapping octadecanol and octadecanal QTL on chromosome 20 in *Heliconius* suggest the presence of a bifunctional FAR that produces both the alcohol and aldehyde together, or alternately tight linkage of separate FAR and alcohol dehydrogenase genes. Further studies, including functional assays and location of wing pheromone biosynthesis, will be required to tease apart our potential candidates.

The presence of a single large-effect QTL for octadecanal production is not surprising, as large-effect loci have been seen in pheromone production in various moth species (Groot et al. 2016, Haynes et al. 2016). What is more surprising is that species differences in moths largely are the result of minor variations in similar compounds or compound ratios, while the production of both octadecanal and its precursor octadecanol is essentially absent in *H. cydno.* Nevertheless, despite the recruitment of stearic acid into this novel product, we see only a single QTL, potentially due to a single gene or tight linkage of two or more biosynthetic genes. The octadecanal locus on chromosome 20 does not overlap with any of the known genes involved in color pattern and mate choice in *Heliconius*, which all lie on other chromosomes (Jiggins 2017), and notably there is no overlap with the *optix* color pattern gene or previously described mate choice QTL to which it is tightly linked (Merrill et al. 2019). Tight linkage of loci for traits under divergent selection and those contributing to premating isolation should facilitate speciation (Felsenstein 1981; Merrill et al. 2010; Smadja and Butlin 2011) but based on our data there is no linkage between olfactory cues and divergent warning patterns in *Heliconius melpomene* and *H. cydno.* It is possible that olfaction does not play a significant role in reproductive isolation. Other color pattern loci are scattered across the *Heliconius* genome, rather than being tightly linked. Instead of acting as a color pattern, mate choice, and pheromone supergene, the loci responsible for these traits are mostly unlinked. Perhaps the selection favoring genetic linkage is weak in these species now that speciation is nearly complete (see also Davey et al. 2017).

Our studies of the electrophysiological and behavioral responses of *Heliconius* butterflies and the genetic basis of pheromone production add to the growing body of literature suggesting that pheromonal communication in Lepidoptera is not limited to nocturnal moths but can be found in day-flying butterflies that also use striking visual signals. *Heliconius* butterflies can detect con- and heterospecific wing compound bouquets, and a major component, octadecanal, is physiologically and behaviorally active in *H. melpomene* and its genetic basis appears relatively simple, consistent with other pheromone shifts found in insects (Symonds and Elgar 2007; Smadja & Butlin 2009). Along with their striking wing color patterns, male *Heliconius* use chemistry to influence female mate choice, combining courtship behaviors, and chemistry in a dance to elicit female mating responses (Klein and de Araújo 2010; Mérot et al. 2015). Despite our human bias towards visual signals, we are now beginning to understand how such visually striking butterflies communicate using chemistry.

## Supporting information

Supplemental results

Sequences of Heliconius cydno candidates

## Author contributions

KJRPB designed, performed, and analyzed electrophysiology experiments, performed QTL mapping and downstream analysis, and wrote the manuscript. KD designed and performed QTL mapping crosses, designed behavioral experiments, and prepared sequencing libraries. KD, SFG, DAA, KJRPB, and JM reared butterflies for all experiments. JM performed and analyzed behavioral experiments. IAW prepared sequencing libraries. YFC and MK provided Tn5 transposase for sequencing. SS provided synthetic compounds used in electrophysiology and behavior experiments as well as mass spectrometry libraries. PR analyzed sequencing data and constructed linkage maps. RMM helped design the QTL experiment. CDJ and WOM contributed funding, resources, and helped design the overall study. All authors contributed to the manuscript editing process.

## Acknowledgments

The authors wish to thank Bill Wcislo and Callum Kingwell for advice on electroantennography. We also thank the insectary team in Panama including Oscar Paneso and Chi-Yun Kuo, as well as the administrative support of Oris Acevedo and Melissa Cano. We also thank S. Dilek for technical assistance. Permits for research and collection of butterfly stocks were provided by the government of Panama. KJRPB, KD, JM, IAW, and CDJ were funded by the European Research Council (FP7-IDEAS-ERC 339873); KD was additionally funded by a Natural Environment Research Council Doctoral Training Partnership and a Smithsonian Tropical Research Institute Short Term Fellowship; YFC and MK are supported by the European Research Council Grant No. 639096 “HybridMiX” and by the Max Planck Society; WOM is funded by the Smithsonian Tropical Research Institute; and SS by the Deutsche Forschungsgemeinschaft (grant DFG Schu984/13-1).

SI Table 1: Comparison of *Heliconius cydno* and *H. melpomene* wing androconia bouquet. Amounts given are the mean in nanograms across 31 *(H. melpomene)* and 26 *(H. cydno)* samples, ± SD. Compounds are only included if found in at least 1/3 of samples for at least one species at levels of at least 1ng. #: compound found in at least 90% of samples of that species. Bold: species differ significantly, dof: Welch’s two sample t-test degrees of freedom.

SI Table 2: Wing region specificity of *Heliconius cydno* androconial compounds. Amounts given are the mean across eight samples ± SD. Numbers in parentheses after compound amounts indicate the number of samples the compound was found in. Bold: wing regions differ significantly. NA: post-hoc test not conducted as original linear model not significant.

SI Table 3: Details of statistical comparisons of stimuli from electrophysiological experiments. P1-P7: as in the main text and Figure 4.

SI Table 4: Short-term adaptation to different stimuli in *H. melpomene* and *H. cydno* males and females, t1, t2, and t3: first, second, and third members of a stimulus triplet. dof: degrees of freedom. P1-P7: as in the main text and Figure 4.

SI Table 5: Potential candidate genes for octadecanal production underlying the QTL peak on chromosome 20 in *Heliconius melpomene.*

SI Figure 1: Comparison of the major compounds in *Heliconius melpomene* and *H. cydno.* Compounds shown are those contributing at least 1% of the total bouquet amount in either species. Numbers under each bar indicate how many samples (out of 31 for *H. melpomene* and 26 for *H. cydno)* the compound was found in. Significantly different compounds: * p < 0.05; ** p < 0.01; *** p < 0.001. n.s., not significant.

SI Figure 2: Absolute and relative abundance of different compound classes in *H. melpomene* and *H. cydno.* n.s., not significant; * p < 0.05; ** p < 0.01; *** p < 0.001. The two unknown categories were not tested as the compound types are not known.

SI Figure 3: The seven compounds found in at least 0.1ng/mm^2^ of wing tissue in at least one wing region in *Heliconius cydno.* A: Compound abundance per square millimeter of tissue. B: Compound abundance without tissue area correction. Numbers under each bar indicate how many samples (out of eight) the compound was found in; letters above bars indicate significant differences between regions, n.s., not significant. A: hindwing androconia; O: forewing overlap region; H: hindwing excluding androconia; F: forewing excluding overlap region.

SI Figure 4: Structures and names of major components of the androconia of *H. melpomene* and *H. cydno* used in electrophysiological experiments.

SI Figure 5: Synthesis of target compounds used in electrophysiological experiments. IBX: iodosobenzoic acid; LiAIH: lithium aluminum hydride.

SI Figure 6: *Heliconius melpomene* responds to electrophysiological stimuli. Top to bottom: dichloromethane plus 2-tetradecyl acetate (internal standard) (negative control), *Lontono* extract (positive control), natural *H. melpomene* male wing extract, depiction of stimulus pulse timing. Data from a single virgin female. Bar-ended lines indicate the measured amplitude of the antennal response.

SI Figure 7: Long-term adaptation to natural and synthetic stimuli in *Heliconius* butterflies. The 95% confidence intervals of the robust LMM slope are shown; a negative slope means that responses to that stimulus drop overtime. P1-P7: see Figure 2.

SI Figure 8: Strength of long-term adaptation correlates with amplitude of EAG response in a sex-specific fashion. In females, a stronger response to a given stimulus correlates with a stronger degree of LTA both overall and for the synthetic compound set. In males the same is seen overall and for the natural extract set in *H. cydno*, but not in *H. melpomene.*

SI Figure 9: Octadecanal persistence in treated males over time. Bars show individual males, with two males per treatment-time point combination, ml-8: separate male individuals.

SI Figure 10: Octadecanal in *H. melpomene, H. cydno*, two Fı families (one in each crossing direction), and the ten backcross to *H. melpomene* families used in QTL mapping. Colors: blue (*H. cydno);* purple (Fı crosses of *H. melpomene* and *H. cydno);* pink (backcrosses to *H. melpomene);* red *{H. melpomene).*

SI Figure 11: Chromosome 20 QTL map for production of octadecanal and octadecanol in *H. melpomene.* Shaded regions indicate the Bayesian confidence intervals with kinship structure taken into account and black line indicates the peak of the QTL.

