## Supplemental figures 1-11 for "A major locus controls a biologically active pheromone component in *Heliconius melpomene*"

SI Figure 1: Comparison of the major compounds in *Heliconius melpomene* and *H. cydno*. Compounds shown are those contributing at least 1% of the total bouquet amount in either species. Numbers under each bar indicate how many samples (out of 31 for *H. melpomene* and 26 for *H. cydno*) the compound was found in. Significantly different compounds: * p < 0.05; ** p < 0.01; *** p < 0.001. n.s., not significant.

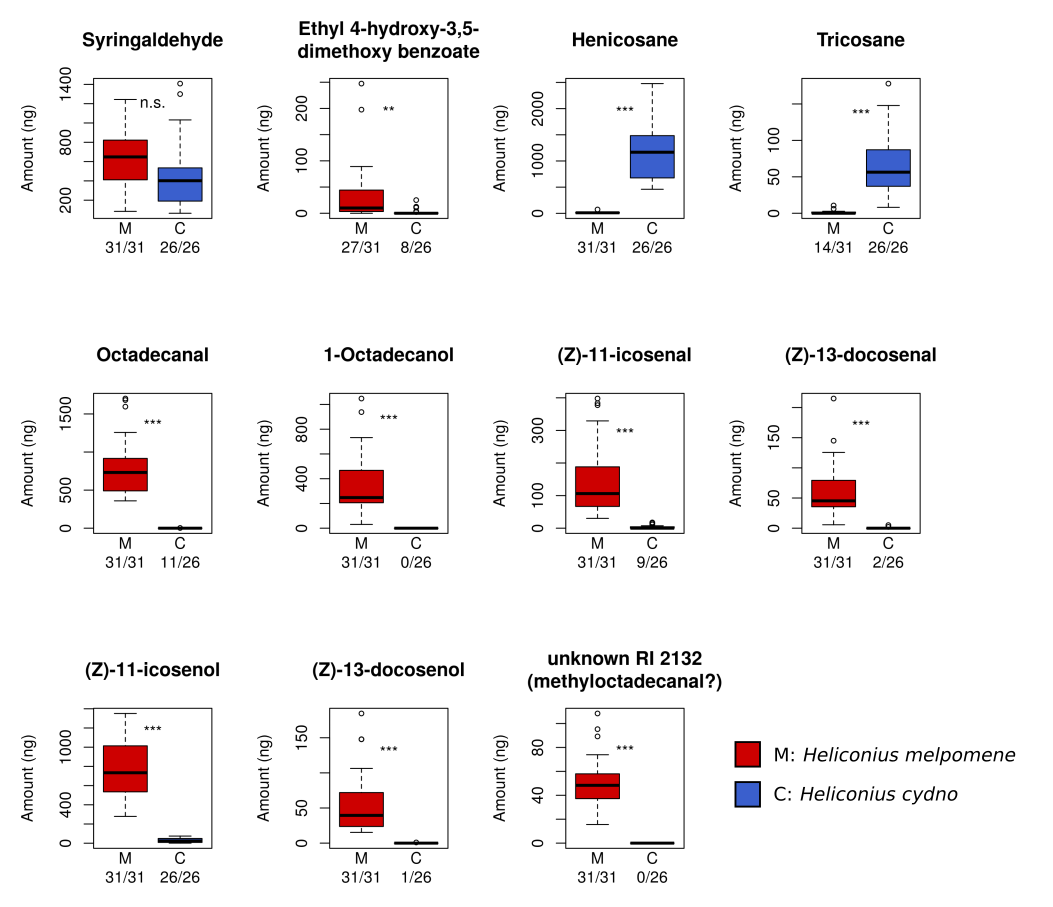

SI Figure 2: Absolute and relative abundance of different compound classes in *H. melpomene* and *H. cydno*. n.s., not significant; * p < 0.05; ** p < 0.01; *** p < 0.001. The two unknown categories were not tested as the compound types are not known.

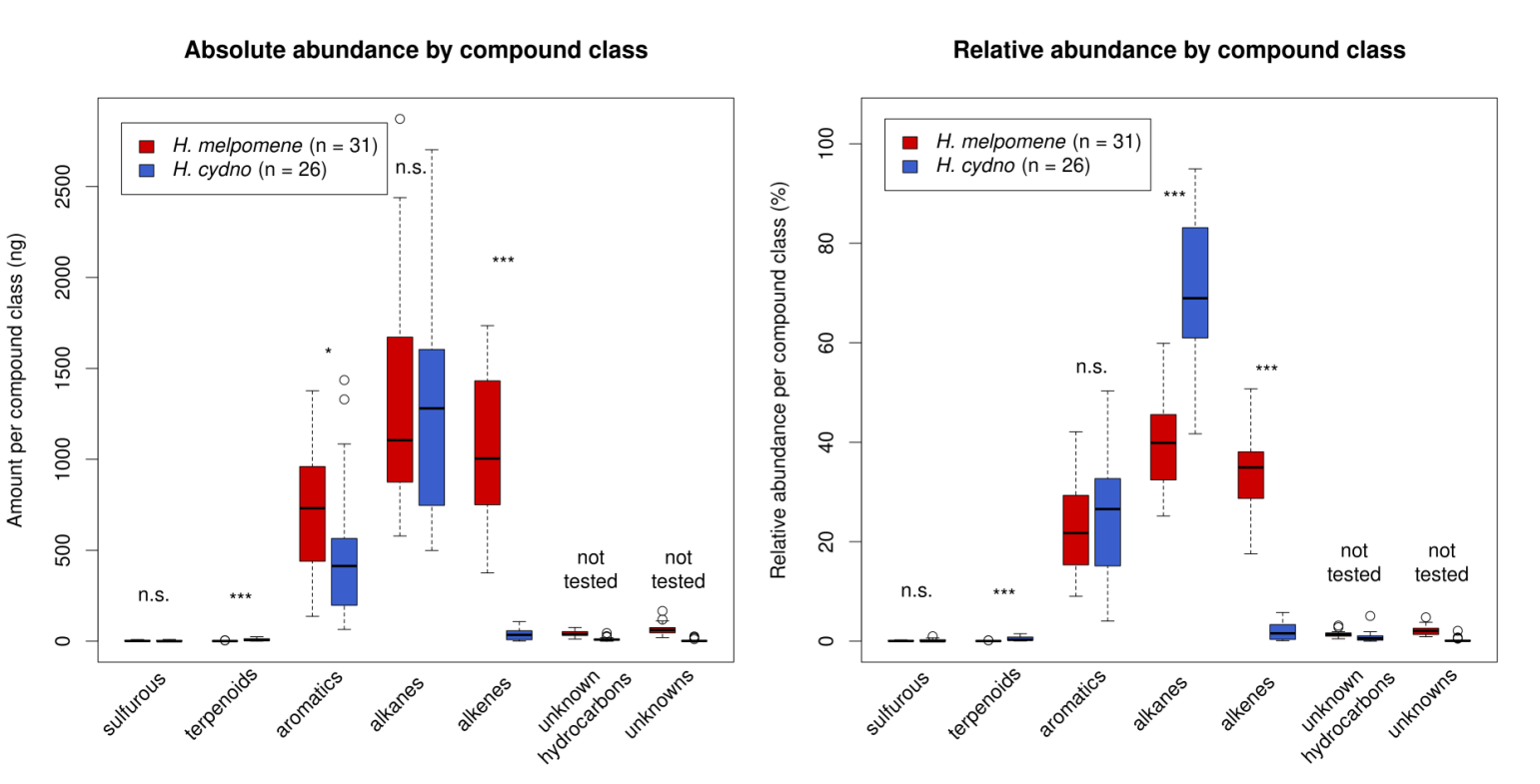

SI Figure 3: The seven compounds found in at least 0.1ng/mm^2^ of wing tissue in at least one wing region in *Heliconius cydno*. A: Compound abundance per square millimeter of tissue. B: Compound abundance without tissue area correction. Numbers under each bar indicate how many samples (out of eight) the compound was found in; letters above bars indicate significant differences between regions. n.s., not significant. A: hindwing androconia; O: forewing overlap region; H: hindwing excluding androconia; F: forewing excluding overlap region.

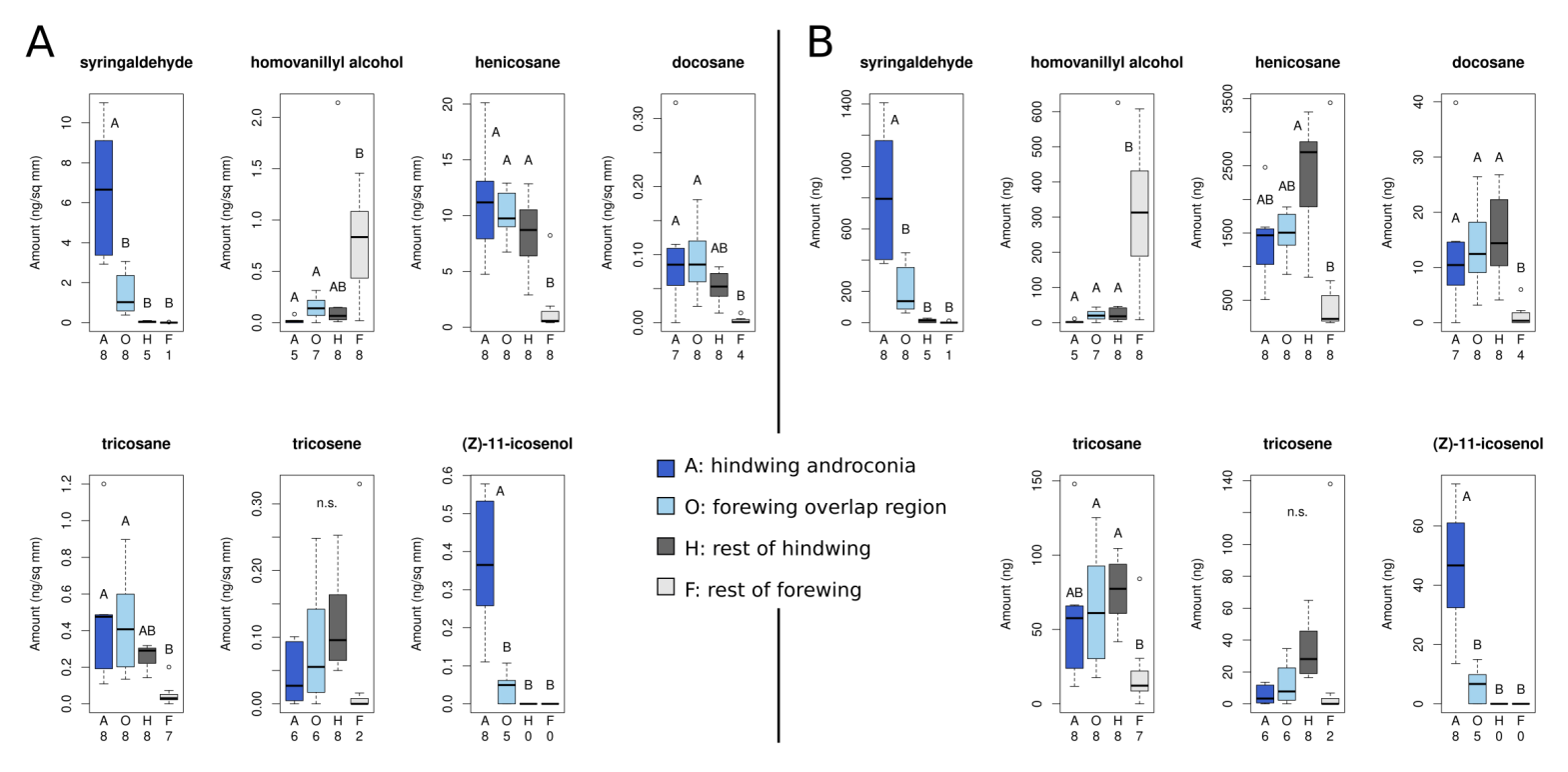

SI Figure 4: Structures and names of major components of the androconia of *H. melpomene* and *H. cydno* used in electrophysiological experiments.

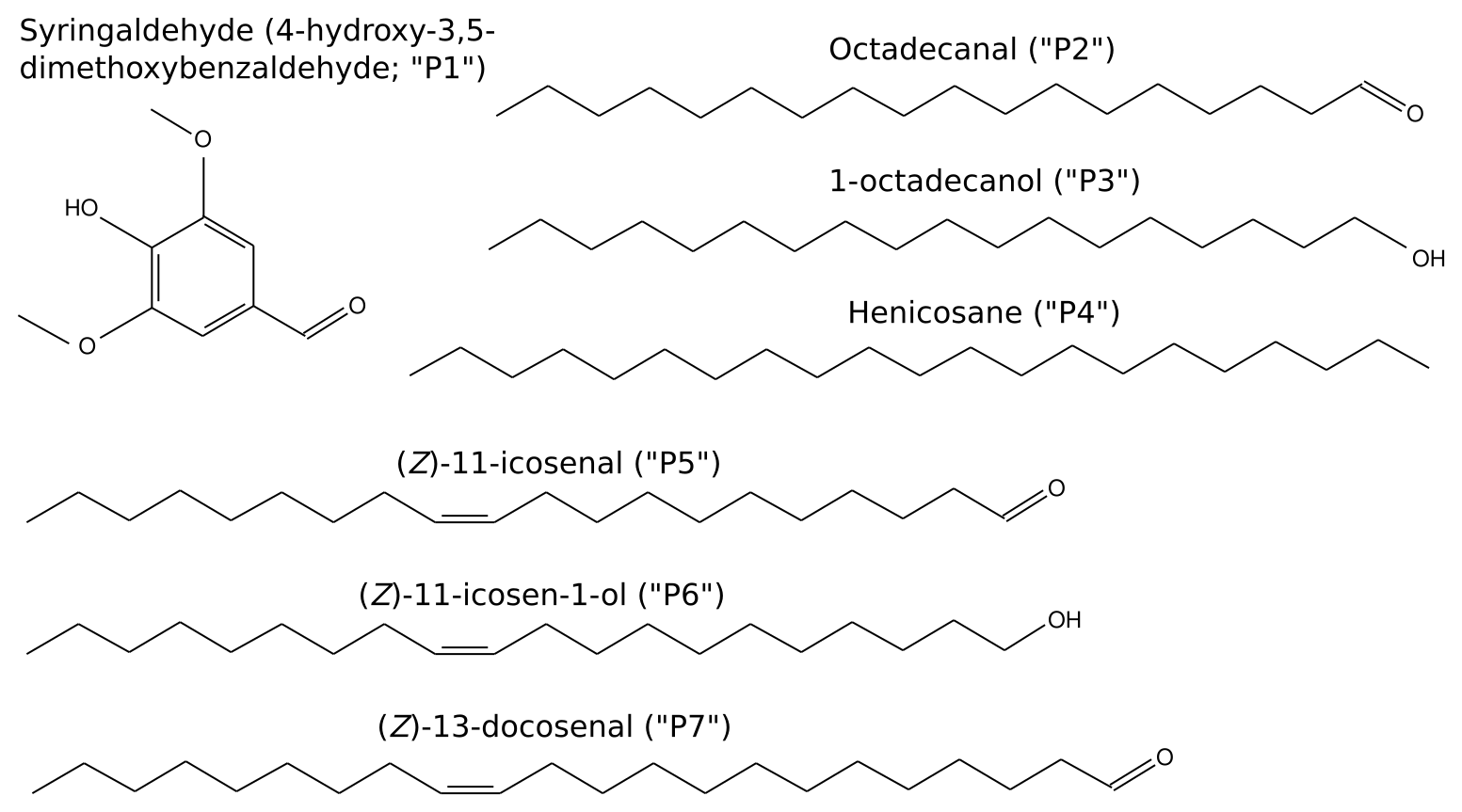

SI Figure 5: Synthesis of target compounds used in electrophysiological experiments. IBX: iodosobenzoic acid; LiAlH: lithium aluminum hydride.

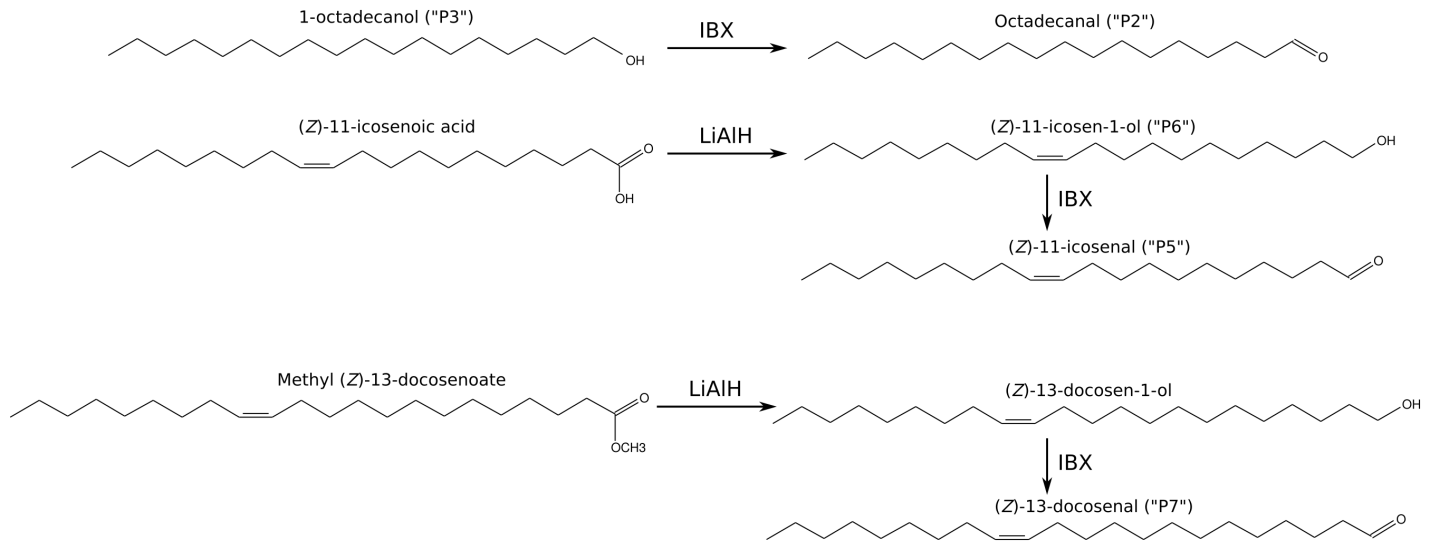

SI Figure 6: *Heliconius* *melpomene* responds to electrophysiological stimuli. Top to bottom: dichloromethane plus 2-tetradecyl acetate (internal standard) (negative control), *Lantana* extract (positive control), natural *H. melpomene* male wing extract, depiction of stimulus pulse timing. Data from a single virgin female. Bar-ended lines indicate the measured amplitude of the antennal response.

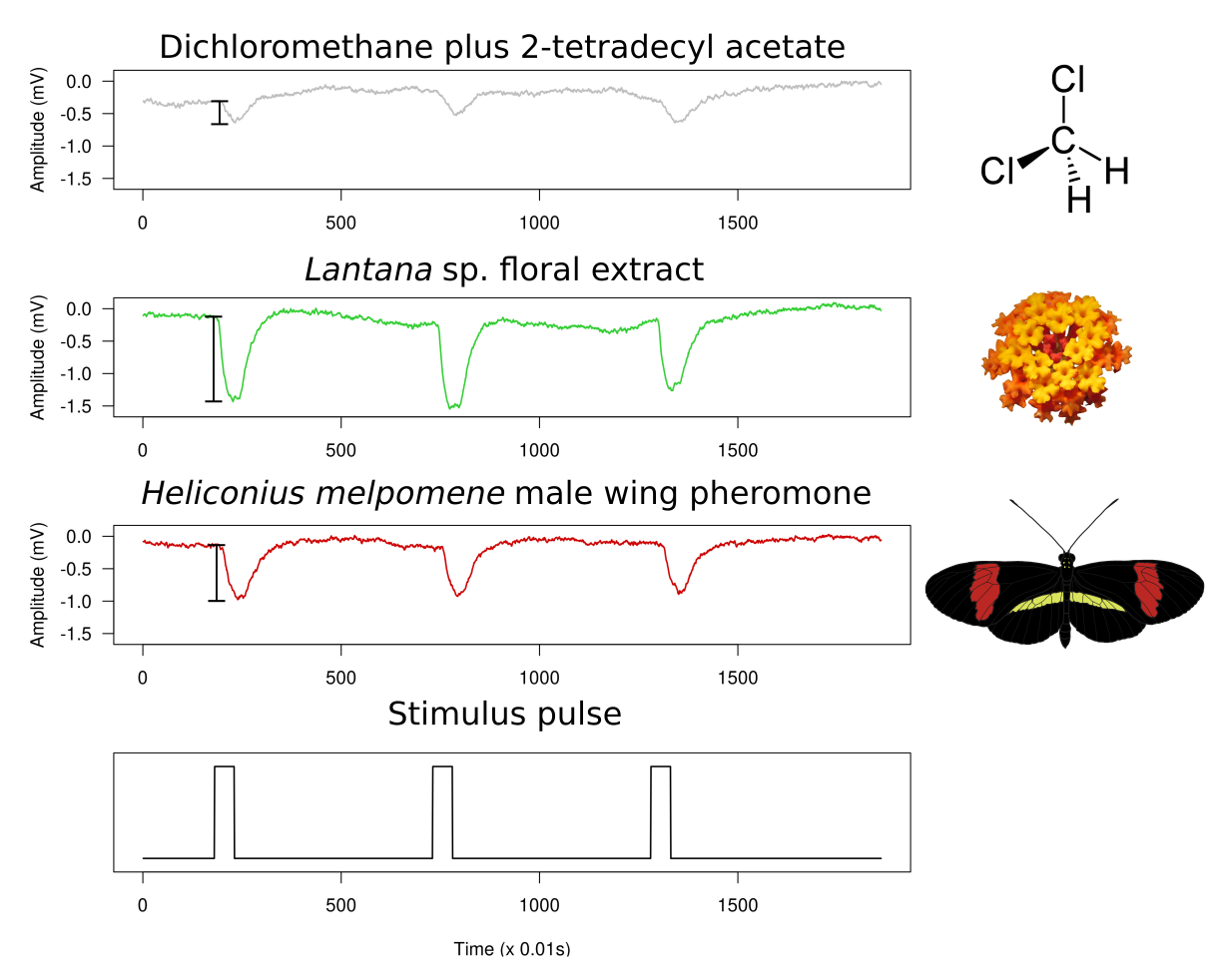

SI Figure 7: Long-term adaptation to natural and synthetic stimuli in *Heliconius* butterflies. The 95% confidence intervals of the robust LMM slope are shown; a negative slope means that responses to that stimulus drop over time. P1-P7: see Figure 2.

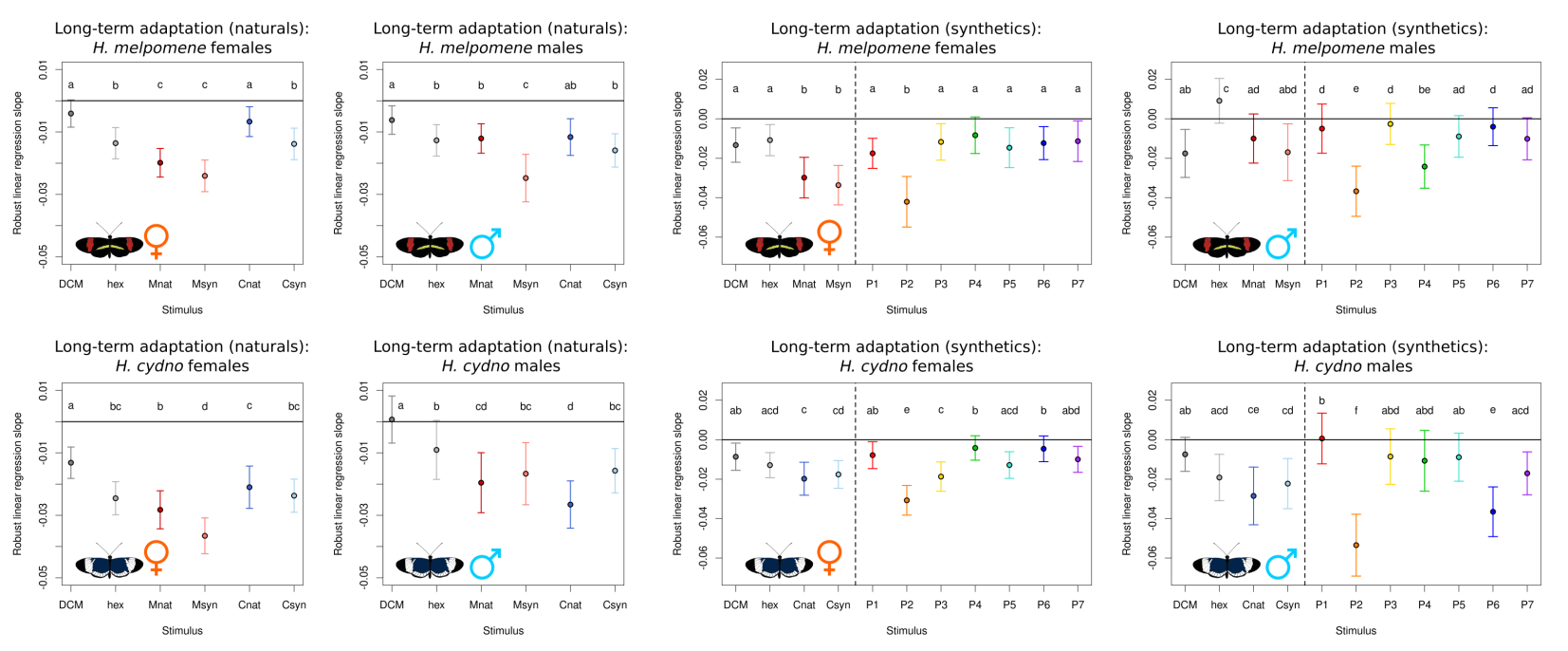

SI Figure 8: Strength of long-term adaptation correlates with amplitude of EAG response in a sex-specific fashion. In females, a stronger response to a given stimulus correlates with a stronger degree of LTA both overall and for the synthetic compound set. In males the same is seen overall and for the natural extract set in *H. cydno*, but not in *H. melpomene*.

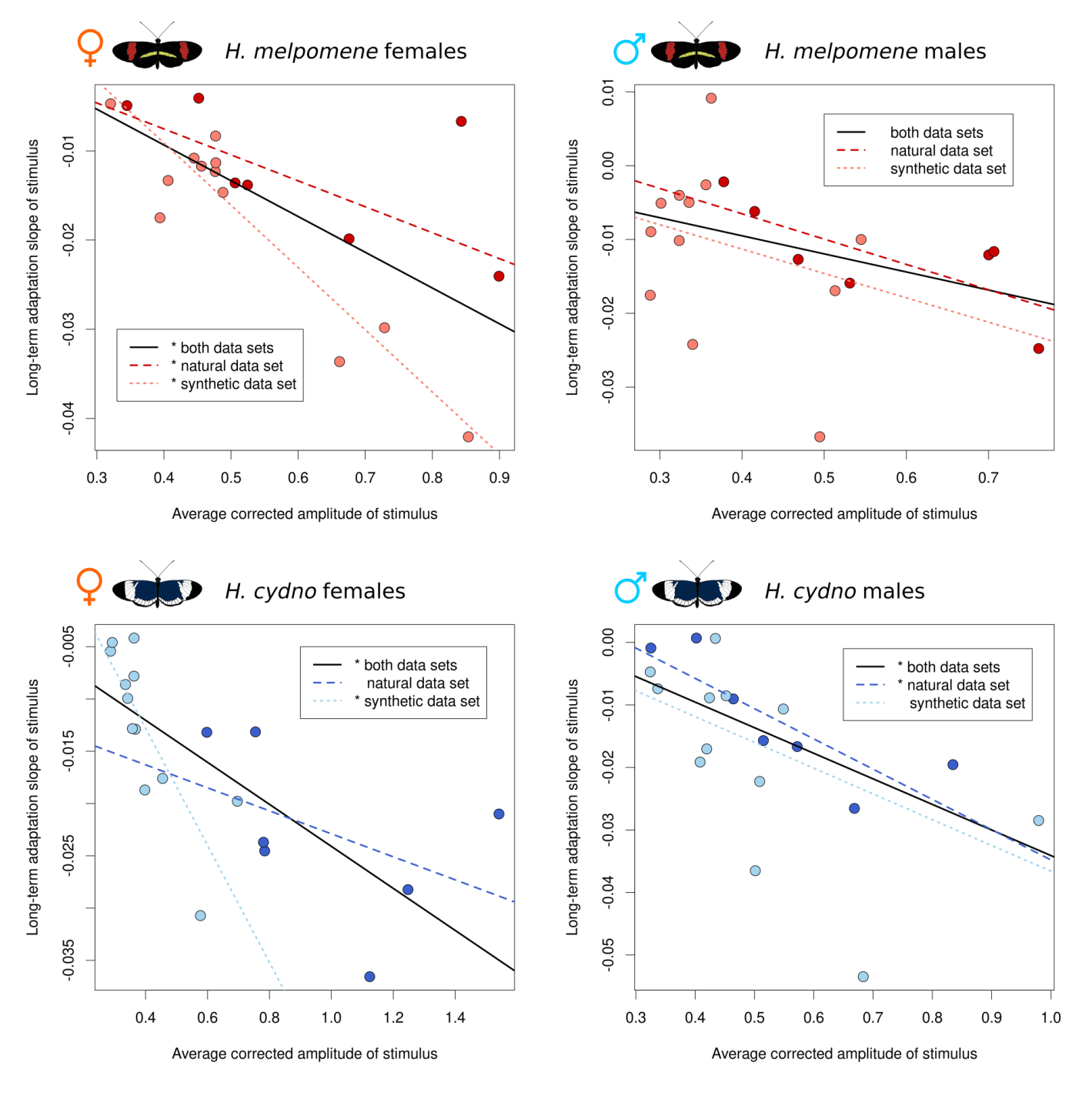

SI Figure 9: Octadecanal persistence in treated males over time. Bars show individual males, with two males per treatment-time point combination. m1-8: separate male individuals.

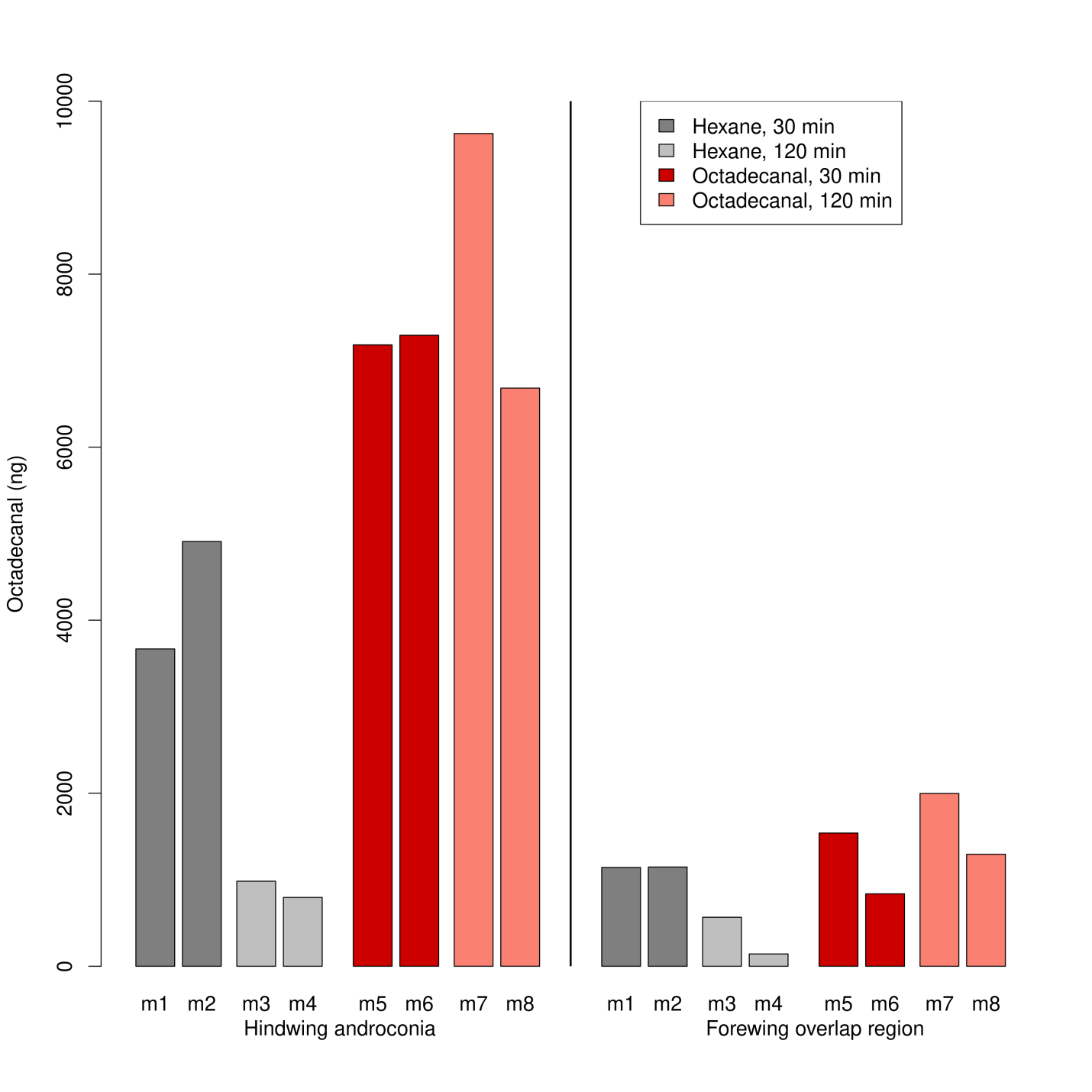

SI Figure 10: Octadecanal in *H. melpomene*, *H. cydno*, two F_1_ families (one in each crossing direction), and the ten backcross to *H. melpomene* families used in QTL mapping. Colors: blue (*H. cydno*); purple (F_1_ crosses of *H. melpomene* and *H. cydno*); pink (backcrosses to *H. melpomene*); red (*H. melpomene*).

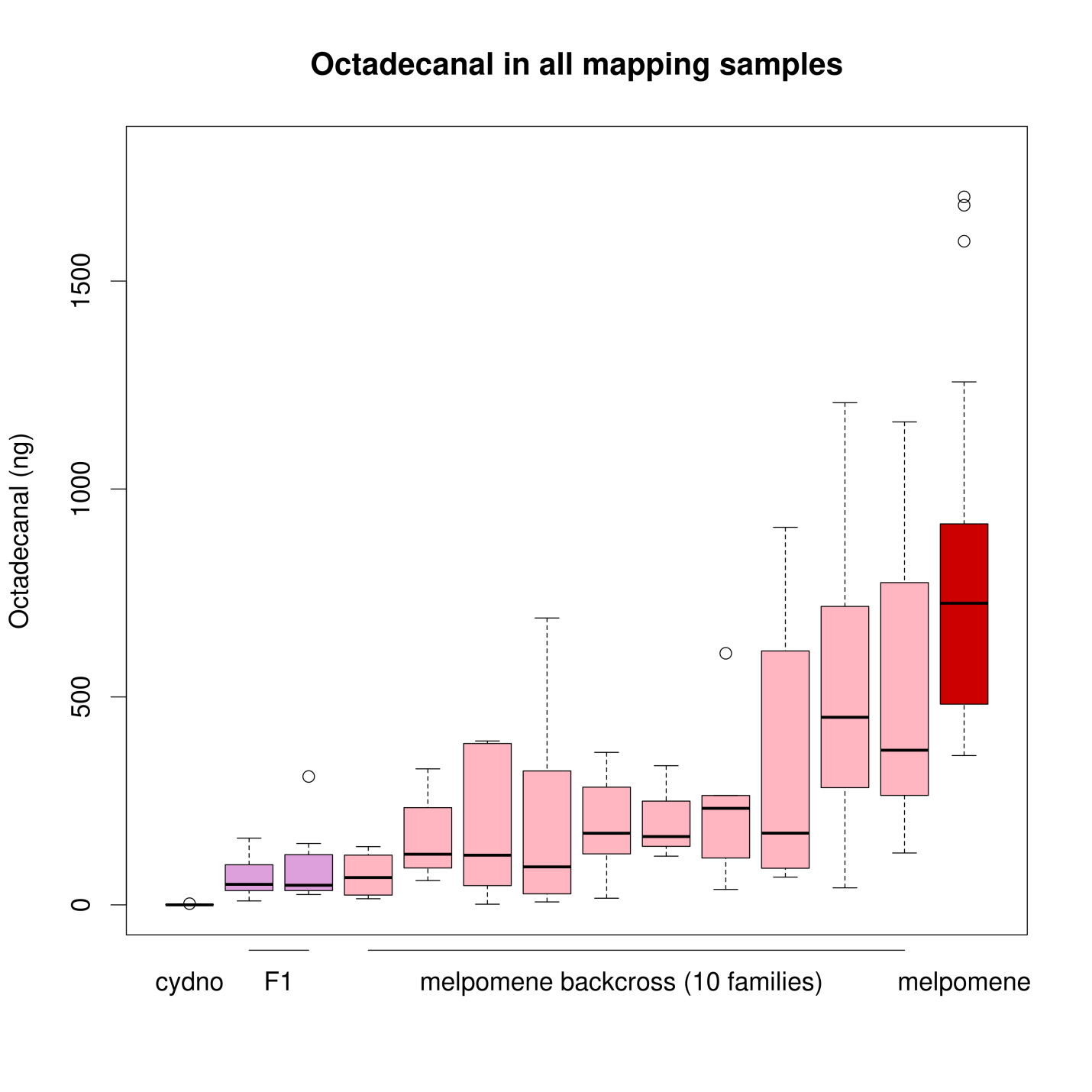

SI Figure 11: Chromosome 20 QTL map for production of octadecanal and octadecanol in *H. melpomene*. Shaded regions indicate the Bayesian confidence intervals with kinship structure taken into account and black line indicates the peak of the QTL.

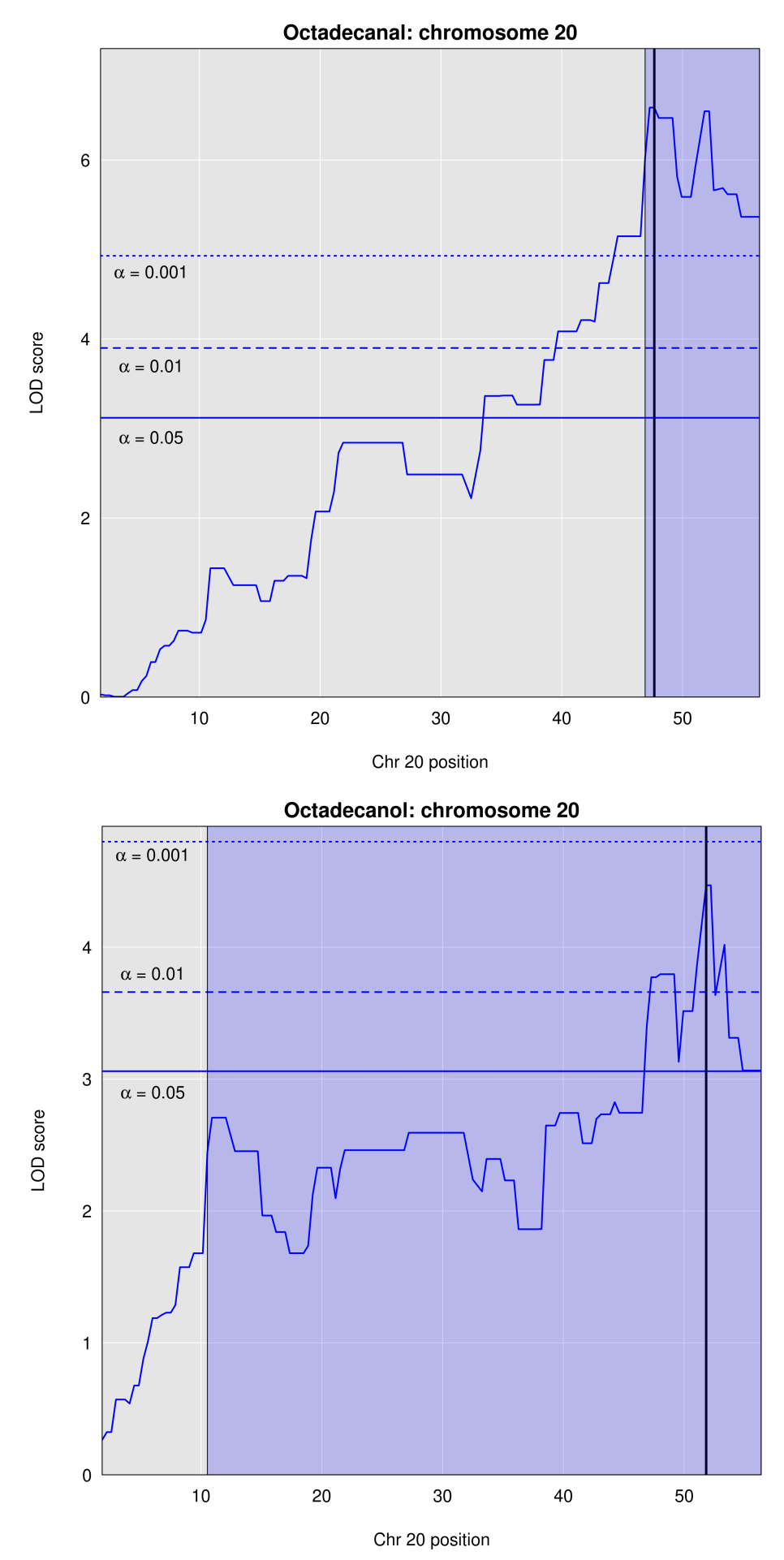
