## Supplemental results for "A major locus controls a biologically active pheromone component in *Heliconius melpomene*"

**SI Results**

Androconial chemical bouquets of *Heliconius melpomene* and *H. cydno*

The analysis of *H. melpomene* included a reanalysis of 18 samples originally published in (Darragh et al. 2017,2019a), as well as 13 previously unpublished samples, for a total of 31. The analysis of *H. cydno* used 26 unpublished samples, eight of which were additionally used for the wing region comparison. Overall, *H. melpomene* produces far more compounds in both amount (3174 ± 1040 ng vs. 1787 ± 776 ng in *H. cydno*, Table S1) and compound diversity (53 compounds found in at least 1/3 of samples vs. 31 compounds in *H. cydno*). Approximately half of all compounds were shared between species (24 of 60 total compounds found in at least 1/3 of samples of both species); of the remainder, the majority of compounds were *H. melpomene*-specific (29 compounds), with a more limited set of *H. cydno*-specific compounds (7 compounds). When grouping compounds by chemical class (Figure S2), *H. melpomene* produced a higher total amount of aromatics and alkenes than *H. cydno*, while *H. cydno* produced more total amount of terpenoids. On a relative basis, the bouquet of *H. melpomene* had a higher percentage of alkenes than *H. cydno*, whereas *H. cydno* was dominated by alkanes. Of the 11 compounds comprising at least 1% of either species’ overall bouquet, all but two (the straight-chain alkanes henicosane and tricosane) were found in equal or higher amounts in *H. melpomene* (Figure S1). In *H. cydno*, two compounds (syringaldehyde and henicosane) together made up more than 90% of the overall compound bouquet.

In *H. melpomene*, several compounds are found exclusively or nearly exclusively in the hindwing androconia region (Darragh et al. 2017). We performed a similar assay using eight of the *H. cydno* samples with all four wing regions (hindwing androconia, forewing overlap region, rest of hindwing, rest of forewing) dissected out. Relative to wing region area, the hindwing androconia produced higher amounts of syringaldehyde and (Z)-11-icosenol than the rest of the wing (Figure S3A), while the other five compounds were found in high amounts in any wing region and were relatively equivalent between the androconia and at least two other wing regions. When compound amounts were not corrected for wing region area, the effects were similar (Figure S3B), except for minor differences between other wing regions. Interestingly, there appeared to be a tradeoff between amounts of syringaldehyde and homovanillyl alcohol across wing regions; as these compounds are structurally similar aromatics this may reflect a shared precursor pool with each wing region being biased towards one or the other product.

Long-term adaptation to pheromone extracts and synthetic compounds

Long-term adaptation (LTA; across the duration of stimulus presentation) was quite common, particularly in responses to natural and synthetic wing pheromone extracts (Figure S7). Adaptation was more common in females, with female *H. melpomene* showing LTA to all stimuli with the exception of henicosane (the only synthetic component tested that is found in low amounts in *H. melpomene*) and disagreement between stimulus sets for air (not shown) and DCM+IS. Female *H. cydno* were similar, with LTA to all stimuli except henicosane, (*Z*)-11-icosenol, and disagreement between stimulus sets for air (not shown). Males showed LTA to fewer stimuli, particularly to synthetic pheromone components; *H. melpomene* males only had LTA to octadecanal and henicosane, and *H. cydno* males only to octadecanal, (*Z*)-11-icosenol, and (*Z*)-13-docosenal. The only natural or synthetic mixture that did not show LTA across all species-sex pairs was *H. melpomene* natural wing (disagreement between stimulus sets in *H. melpomene* males). For synthetic pheromone components, the only disagreements between species within a sex were in (*Z*)-11-icosenol (for females) and henicosane, (*Z*)-11-icosenol, and (*Z*)-13-docosenal (for males).

Long-term adaptation was often stronger to natural stimuli when compared with the two solvents (hexane and DCM+IS). In *H. melpomene* females, adaptation to *H. melpomene* natural and synthetic wing pheromone was stronger than adaptation to DCM+IS, while adaptation to both natural and synthetic *H. cydno* wing pheromone was equivalent to the relevant solvent (DCM+IS and hexane respectively). The same pattern was found in *H. melpomene* males. In *H. cydno* females, only synthetic *H. cydno* wing pheromone showed equivalent LTA to its hexane solvent, while LTA to all other stimuli was stronger than their respective solvents. The pattern differed in *H. cydno* males, with only natural wing extracts from both species having stronger LTA than their solvent (DCM+IS). By contrast with natural stimuli, LTA to pheromone components was often equivalent to the two solvents. In *H. melpomene* females, only octadecanal differed from the hexane solvent, and the strength of LTA to octadecanal was identical to that of natural and synthetic *H. melpomene* wing pheromone. Other species-sex combinations showed no obvious pattern in comparison to the solvents, although both sexes of *H. cydno* showed stronger LTA to *H. cydno* wing pheromone than to its DCM+IS solvent.

In general, LTA to *H. melpomene* pheromones was stronger than to *H. cydno* pheromones. Females of both species had different degrees of LTA to the two species’ wing pheromones (in both cases, LTA was stronger to *H. melpomene* wings), while males had an equal degree of LTA. There was no obvious link between the presence or absence of LTA to a given pheromone component and the presence or absence of a given pheromone component in the preparation’s species’ wing pheromones. A significant correlation existed between degree of LTA and strength of response in to a given stimulus (Figure S8). Females of both species showed significant correlations for the synthetic stimulus sets and the overall data, while males of both species showed significant correlations between LTA strength and response strength for the natural stimulus sets and the overall data.

Short-term adaptation to pheromone extracts and synthetic compounds

By contrast to LTA, short-term adaptation (STA) was mostly absent in all combinations of preparation species-sex and stimulus type (Table S4). No stimuli showed STA across all species-sex preparation combinations. Both female and male *H. melpomene* had STA to *H. cydno* wing pheromones. Female *H. cydno* also showed STA to *H. melpomene* wing extract. Female *H. melpomene* showed STA to 1-octadecanol, while male *H. cydno* showed STA to 1-octadecanol and (*Z*)-11-icosenal.
